# Evolution of cation binding in the active sites of P-loop nucleoside triphosphatases

**DOI:** 10.1101/420133

**Authors:** Daria N. Shalaeva, Dmitry A. Cherepanov, Michael Y. Galperin, Armen Y. Mulkidjanian

**Affiliations:** School of Physics, University of Osnabrück, D-49069 Osnabrück, Germany; A.N. Belozersky Institute of Physico-Chemical Biology, Lomonosov Moscow State University, Moscow 119992, Russia; School of Bioengineering and Bioinformatics, Lomonosov Moscow State University, Moscow 119992, Russia; Semenov Institute of Chemical Physics, Russian Academy of Sciences, Moscow 119991, Russia; National Center for Biotechnology Information, National Library of Medicine, National Institutes of Health, Bethesda, Maryland 20894, USA

## Abstract

The activity of cellular nucleoside triphosphatases (NTPases) must be tightly controlled to prevent spontaneous ATP hydrolysis leading to cell death. While most P-loop NTPases require activation by arginine or lysine fingers, some of the apparently ancestral ones are, instead, activated by potassium ions, but not by sodium ions. We combined comparative structure analysis of P-loop NTPases of various classes with molecular dynamics (MD) simulations of Mg-ATP complexes in water and in the presence of potassium, sodium, or ammonium ions. In all analyzed structures, the conserved P-loop motif keeps the triphosphate chains of enzyme-bound NTPs in an extended, catalytically prone conformation, similar to that attained by ATP in water in the presence of potassium or ammonium ions bound between alpha- and gamma-phosphate groups. The smaller sodium ions could not reach both alpha- and gamma-phosphates of a protein-bound extended phosphate chain and therefore are unable to activate most potassium-dependent P-loop NTPases.

## Introduction

P-loop nucleoside triphosphatases (NTPases) represent the most common protein fold that can comprise up to 18% of all gene products in a cell (1–4). P-loop NTPase domains, which apparently preceded the Last Universal Cellular Ancestor (4–10), are found in translation factors, small GTPases, kinases, helicases, rotary ATP synthases, and many other ubiquitous proteins.

The P-loop fold, a variation of the Rossman fold, is a 3-layer αβα sandwich, where the N-terminal β- strand is connected with the following α-helix by an elongated flexible loop typically containing the GxxxxGK[ST] sequence motif, known as the Walker A motif (11), see Fig. 1. This motif is responsible for binding the NTP’s phosphate chain and is often referred to as the P-loop (*<u>p</u>*hosphate-binding loop) motif (12). The conserved lysine residue of the P-loop forms hydrogen bonds (H-bonds) with β- and γ-phosphate groups, while the following Ser/Thr residue coordinates the Mg^2+^ ion, which, in turn, coordinates β- and γ-phosphates from the other side of the phosphate chain (Fig. 1A-C). Another motif typical for P-loop proteins is the Walker B motif with the sequence pattern *hhhh*D, where “*h*” denotes a hydrophobic residue (11). In P-loop NTPases, the aspartate from this motif either serves as a direct Mg^2+^ ligand or participates in the second coordination sphere of Mg^2+^. Further specific motifs are shown in Fig. 1.

**Figure 1.**
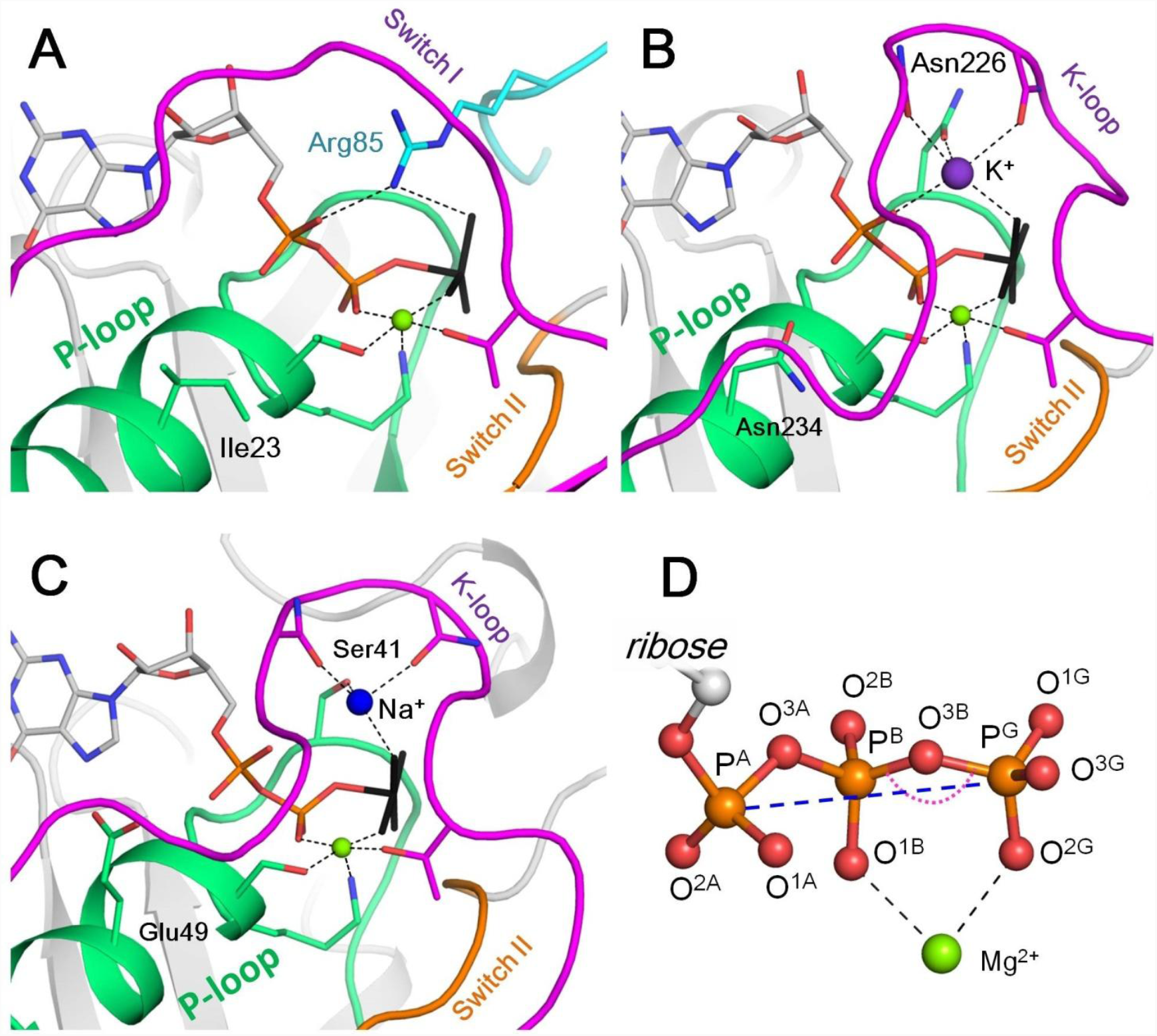
Mg-NTP complexes and their binding in the active sites of P-loop NTPases. Phosphate chains of NTP molecules and their analogs are colored by atoms: oxygen atoms in red, phosphorus in orange. The K^+^ ion is shown as a purple sphere, Na^+^ ion is shown as a blue sphere, Mg^2+^ ions are shown as green spheres. Phosphate chain is shown in stick representation with oxygens in red and phosphorus atoms in orange; γ-phosphate mimicking groups (AlF_4_^-^ and MgF_3_) are shown in black, coordination and hydrogen bonds are shown as black dashed lines. **A**. Active site of the small Ras-like GTPase RhoA in complex with the activating protein RhoGAP [PDB entry 1OW3]; the bound GDP-MgF_3_ mimics the transition state. The P-loop with the preceding α-helix is shown as green cartoon; Switch I motif with the conserved Mg^2+^-binding Thr residue is shown in magenta; Switch II motif (DxxG motif, which starts from the conserved Asp of the Walker B motif) is shown in orange; the Arg finger of RhoGAP is colored turquoise. **B**. Active site of the K^+^-dependent GTPase MnmE with bound GDP-AlF_4_ [PDB: 2GJ8]. Switch I region and the K-loop are shown in magenta. **C**. The active site of dynamin, a Na^+^-adapted GTPase with bound GDP-AlF_4_ [PDB: 2X2E]. The P-loop and K-loop (Switch I region) are colored as in panels A and B. **D.** Structure of the NTP triphosphate chain with Mg^2+^ ion in a bidentate coordination, referred to as the βγ conformation. The pink dotted arch indicates the P^B^-O^3B^-P^G^ angle; the blue dashed line indicates the P^A^-P^G^ distance. The atom names are in accordance with the CHARMM naming scheme (66) and the recent IUPAC recommendations (67).

Catalytic activity of P-loop NTPases usually requires interaction with other proteins or domains that insert activating Arg or Lys “fingers” in the catalytic site (13), see Fig. 1A. Some P-loop NTPases, instead, functionally depend on monovalent cations (14–20) (Fig. 1B, C, Table S1). Strict dependence on K^+^ ions was shown, among others, for the bacterial tRNA-modifying GTPase MnmE (also known as TrmE), Era-like GTPase Der from *Escherichia coli, AtNOS/AtNOA1* GTPase from *Arabidopsis thaliana*, ribosome assembly GTPase YqeH, G-protein coupled with ferrous transporter FeoB, ribosome-binding ATPase YchF, bacterial ribosome biogenesis protein RgbA, and the DNA repair and recombination protein Rad51 (15–18, 21–29). The requirement for K^+^ or NH_4_^+^ ions was shown both for the intrinsic and ribosome-dependent GTPase activity of several ubiquitous translation factors (30–37). Based on the K^+^-dependence of several ancient ATPases and GTPases of the TRAFAC class, we have previously suggested that it was an ancestral trait which was subsequently replaced by reliance on arginine or lysine fingers (38, 39).

In K^+^-dependent P-loop NTPases, the catalytically important K^+^ ion occupies the position of the positively charged nitrogen atom of the Arg/Lys finger, interacting with the phosphate groups of the NTP molecule from the opposite side of the Mg^2+^ ion (15, 20), see also Fig. 1. Using crystal structures of several related K^+^-dependent P-loop NTPases, a set of their characteristic features could be identified, including a specific K^+^-binding “K-loop” and two specific Asn/Asp residues in the P-loop (18, 20, 38) (Fig. 1B, C). Still, the molecular mechanism(s) of activation of P-loop NTPases either by Arg or Lys fingers or by monovalent cations (hereafter M^+^ ions) remain unresolved, see (40–44) for recent reviews.

In the majority of K^+^-dependent P-loop NTPases, Na^+^ ions could not replace K^+^ ions as cofactors (17, 21–23, 25, 26). The very existence of ubiquitous K^+^-dependent NTPases, along with the strict dependence of the translation system on cytoplasmic K^+^ ions and its inhibition by Na^+^ ions (32), require maintaining the [K^+^]/[Na^+^] ratio >> 1.0 in the cytoplasm. Since Na^+^ usually prevails over K^+^ in natural habitats, cells may spend up to a half of the available energy to maintain the proper [K^+^]/[Na^+^] ratio (45). It has been argued that the first cells emerged in K^+^-rich environments, which could explain the K^+^ dependence of the evolutionarily old cell processes (38). However, it has remained obscure why, in the course of evolution, the cellular machinery has not switched its specificity from K^+^ to Na^+^, considering the obvious similarity of K^+^ and Na^+^ ions and the abundance of Na^+^ in natural habitats (46). Such adaptation would have been widely beneficial, especially in the case of marine organisms, which invest large efforts into counteracting the [K^+^]/[Na^+^] ratio of ∼0.02 in the sea water (47). For P-loop NTPases, the use of Na^+^ ion as an activating cofactor is, in principle, possible: human dynamin and the dynamin-like protein from *A. thaliana* are equally well activated by Na^+^ and K^+^ ions (48, 49). The structures of dynamins show that Na^+^ ions bind in a similar position to that occupied by K^+^ ions in potassium-dependent NTPases (20), *cf*. Fig. 1B and 1C. Therefore, the strong preference of other NTPases for K^+^ ions remains a mystery.

The specific role of K^+^ ions in processing phosphoanhydride bonds has been documented also in the absence of enzymes. Back in 1960, larger ions, such as K^+^ and Rb^+^, were shown to be more efficient than the smaller Na^+^ and Li^+^ ions in accelerating transphosphorylation (50), see Table S2. These observations suggested that the observed catalytic effect of the positive charges of Arg/Lys fingers or K^+^ ions could be determined by the size of these cations.

So far, computational studies of the mechanisms of NTP hydrolysis in water have been conducted using such model systems as methyl triphosphate molecule with and without Mg^2+^, Mg-ATP complex, and Mg-GTP complex, see e.g. (51–55). To our knowledge, no computational studies of Mg-NTP complexes investigated the effects of monovalent cations.

Here, we performed evolutionary analysis of the conformations of NTPs and their analogs bound in the active sites of different families of P-loop NTPases and complemented this analysis with molecular dynamics (MD) simulations of the Mg-ATP complex in water in the presence of K^+^, Na^+^, and NH_4_^+^ ions. We report that, in MD simulations, M^+^ ions got bound to the phosphate chain in the same two sites that are taken by positive charges in the active sites of P-loop NTPases, namely, between β- and γ-phosphates, in the position of the amino group of the invariant P-loop lysine, and between α- and γ-phosphates, in the position that is occupied either by the side chain of the activating Arg/Lys finger or by an M^+^ ion. However, the extended conformation of the phosphate chain, which is similar to the catalytically prone conformation of tightly bound Mg-ATP complexes in the active sites of P-loop NTPases, was achieved only in the presence of the larger K^+^ and NH_4_^+^ ions, but not with the smaller Na^+^ ions. In addition, the comparative structural analysis has revealed that although the activating M^+^ ions are bound exclusively by the residues of the P-loop NTPase domain, the activation of respective NTPases additionally requires a specific interaction of the P-loop domain with the respective activating moiety (another protein domain or an RNA/DNA molecule) to shape the cation-binding site. Such a mechanism prevents uncontrolled hydrolysis of the cellular ATP stock, which, otherwise, could cause cell death.

## Results

### Cation binding to the Mg^2+^-ATP complex

We have conducted a series of molecular dynamics (MD) simulations of the Mg^2+^-ATP complex in water and in the presence of K^+^, Na^+^, or NH_4_^+^ ions (see Methods and Table S3 for details).

As a starting point for the MD simulations, we chose the conformation of Mg-ATP complex with the Mg^2+^ ion coordinated by two non-bridging oxygen atoms of the β- and γ-phosphate groups and four water molecules (Fig. 1D). This mode of Mg^2+^ coordination, often referred to as bidentate or βγ coordination, has been observed in NMR studies of the Mg-ATP complex in water (56–59) and in crystal structures of P-loop NTPases with bound NTPs and their analogs (11, 12, 60–62), see also Fig. 1. The initial structure of the Mg-ATP complex was optimized in vacuum using the PM3 Hamiltonian. After that, 1,200 water molecules and 6 monovalent cations (K^+^, Na^+^ or NH_4_^+^) were added to the Mg-ATP complex. In each case, 4 Cl^-^ ions were added to balance the total charge of the simulation system. The resulting solution corresponded to the total ionic strength of 0.2 M. To investigate the conformational space of the Mg-ATP complex in water, we performed three independent MD simulation runs of 170 ns for each system. During each simulation, the system coordinates were saved every 50 picoseconds, providing 10,000 conformational states (frames) for each system. For the visualization in Fig. 2A, we have selected every 100th simulation frame to sample the conformational states of the Mg-ATP complex with 5-ns intervals. The conformations were superposed to achieve the best possible match between coordinates of the phosphorus and ester oxygen atoms of the ATP phosphate chain.

**Figure 2.**
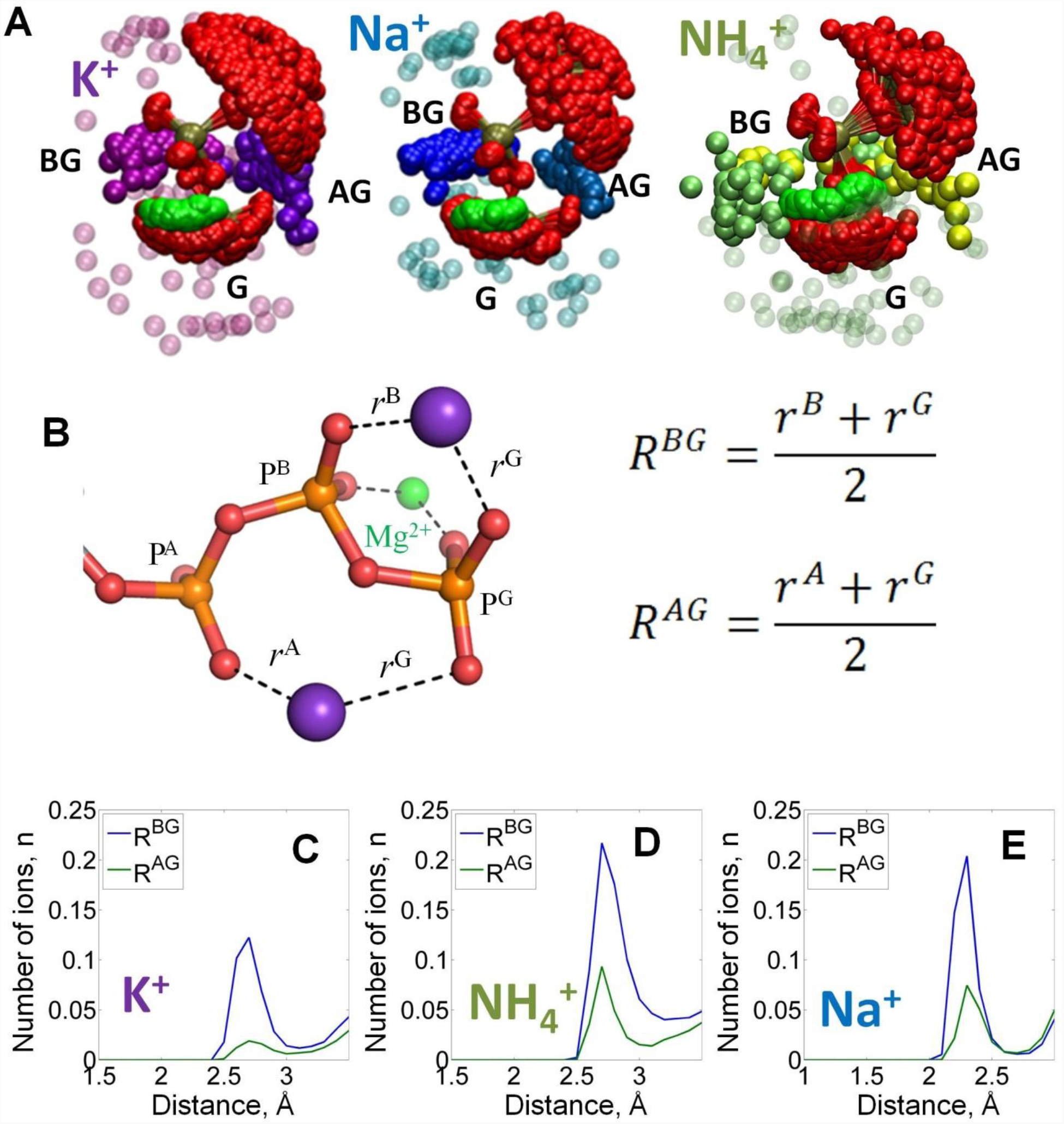
Binding of monovalent cations to the Mg-ATP in water. The color scheme is as in Fig. 1. A, Superposition of the ATP phosphate chain conformations observed in the MD simulations in the presence of K^+^ ions (shown in purple); Na^+^ ions (shown in blue) and NH_4_^+^ ions (nitrogen atoms of NH_4_^+^ ions are shown in yellow/green). The ribose and adenine moieties are not shown, the phosphate chain is shown with P^A^ on top and P^G^ at the bottom. All cations within 5Å from the phosphate chain are shown and colored in different shades depending on the nearby oxygen atoms to illustrate the distinction between binding in the AG and BG sites (see text for details). Transparent spheres signify the ions outside the AG and BG sites. The constellation of ions in the vicinity of γ- phosphate is referred to as the site G. B, Geometry of the Mg-ATP complex with two monovalent cations bound, one in the AG site and one in the BG site. Distances to the AG and BG binding sites (R^AG^ and R^BG^) were calculated as averages of the distances to the two corresponding oxygen atoms. The distances to the oxygen atoms (e.g. r^A^) were defined as the shortest distances between a particular M^+^ ion and any oxygen atom of the respective phosphate group (including ester oxygen atoms). C-E, distance distributions for K^+^, NH_4_^+^, and Na^+^ ions in the AG and BG sites.

Distance distributions obtained from the MD simulation data (Fig. S1) show that M^+^ ions formed coordination bonds with oxygen atoms of the ATP phosphate chain with the respective lengths of 2.2 Å for Na^+^, 2.6 Å for K^+^, and 2.7 Å for NH_4_^+^ ions. These distances correspond well with the crystallographic data for these ions (63–65). On time average, within the 4 Å radius around the phosphate chain, 1.5 cations were present in the case of Na^+^ and NH_4_^+^, and 0.75 cations were present in the case of K^+^ (Fig. S2). Based on the radial distributions of M^+^ ions around each individual oxygen atom of the ATP phosphate chain (Fig. S1) and visual inspection of the M^+^ binding to the phosphate groups, at least two distinct binding sites for M^+^ ions could be identified (Fig. 2A). One of them was formed by the oxygen atoms of β- and γ-phosphates, and the other site involved the oxygens of α- and γ-phosphates. We refer to these binding sites as the BG and AG sites, respectively. Additionally, M^+^ ions were often found close to the distal end of the phosphate chain, where they contacted one or more oxygen atoms of the γ-phosphate (the G site(s), Fig. 2A).

To characterize M^+^ binding in the AG and BG sites, we measured the distances from each M^+^ ion to the nearest oxygen atoms of the two respective phosphate residues (R^AG^ and R^BG^ distances in Fig. 2B). Site occupancy was estimated, as shown in Fig. 2C-E, from the number of M^+^ ions located in the proximity of the binding site at each moment of the simulation. In the BG site, binding of any M^+^ ion produced a prominent maximum in the R^BG^ distribution. The R^BG^ values peaked at the same distance as the maxima of the distribution of distances to separate oxygens (Fig. S1), which indicates that the cations in the BG site simultaneously formed coordination bonds with two oxygen atoms. Similarly, in the AG site, the NH_4_^+^ and Na^+^ ions produced peaks in the R^AG^ distribution plots with the maxima at 2.7 Å and 2.3 Å, respectively. For K^+^ ions, the corresponding peak with a R^AG^ value of 2.6 Å was wide. Still, the distributions of the distances between cations and individual oxygen atoms of the triphosphate chain show that α- and γ-phosphates had the most contacts with K^+^ ions, see graphs for O^2A^ and O^1G^ in Fig. S1 (hereafter, the atom names follow the CHARMM naming scheme (66) and the recent IUPAC Recommendations (67), as shown in Fig. 1D and Fig. S1).

While occupying the same binding sites, M^+^ ions bound with different affinity that decreased in the order of Na^+^ > NH_4_^+^ > K^+^ (Table S2). Higher affinity of ATP to Na^+^ ions, as compared to K^+^ and NH_4_^+^ ions, was previously observed in several experimental studies, albeit in the absence of Mg^2+^ (Table S2). For each M^+^ ion, MD simulation data indicated much lower occupancy of the AG site than of the BG site; the average occupancy of the BG site was estimated to be 0.95 for Na^+^, 0.72 for NH_4_^+^, and 0.5 for K^+^, compared to the average occupancy of the AG site of 0.15 for Na^+^, 0.2 for NH_4_^+^, and 0.05 for K^+^ (Fig. 2C-E).

The reasons for the weak K^+^-binding in the AG site could be, in principle, clarified by structural and thermodynamic analysis of the conformations of the Mg-ATP complex with two K^+^ ions bound. Such an analysis, however, was hindered by the scarcity of the respective MD simulation frames. Therefore, we have conducted additional MD simulations with positional restraints applied to the cations. We have conducted 10-ns simulations of an ATP molecule with Mg^2+^ in the βγ coordination and K^+^ in the BG site, and of the same system but with the addition of the second K^+^ ion in the AG site. Positional restrains were applied to K^+^ and Mg^2+^ ions and to one of the atoms of the adenine base. Binding of the second K^+^ ion in the AG site was found to stabilize all three phosphate groups in a near-eclipsed conformation, with the phosphorus-oxygen bonds of the α-phosphate group almost coplanar to the respective bonds of β- and γ-phosphates (Fig. S3, Table S4). In this conformation, the distance between the oxygen atoms of α- and γ-phosphates was short enough to accommodate the second K^+^ ion. As shown in Fig. S3, binding of the second K^+^ ion in the AG site promotes the transition of the phosphate chain into the almost fully eclipsed conformation by approximately 60 meV or 5.7 kJ/mol.

We were mostly interested in the βγ conformations of the Mg-ATP complex that are typical for P-loop NTPases. To sample enough βγ conformations, we have conducted an additional series of 25 independent 20-ns long MD simulations, with and without M^+^ ions (Table 1). These data were used to define the shape of the phosphate chain of the βγ-coordinated Mg-ATP complex.

**Table 1.** Effects of monovalent cations on the shape of the triphosphate chain of the Mg^2+^-ATP complex in water, as inferred from the MD simulation data.

| Added cation | Conformation of the triphosphate chain of $\text{Mg}$-ATP<sup>a</sup> | | | | | |
| --- | --- | --- | --- | --- | --- | --- |
| | $\beta\gamma$-coordination | | $\beta\gamma$-coordination, "curled" phosphate chain | | $\alpha\beta\gamma$-coordination | |
| | $\text{P}^{\text{A}}\text{-P}^{\text{G}}$<br>distance, Å | $\text{P}^{\text{B}}\text{-O}^{3\text{B}}\text{-P}^{\text{G}}$<br>angle | $\text{P}^{\text{A}}\text{-P}^{\text{G}}$<br>distance, Å | $\text{P}^{\text{B}}\text{-O}^{3\text{B}}\text{-P}^{\text{G}}$<br>angle | $\text{P}^{\text{A}}\text{-P}^{\text{G}}$<br>distance, Å | $\text{P}^{\text{B}}\text{-O}^{3\text{B}}\text{-P}^{\text{G}}$<br>angle |
| None | $5.46 \pm 0.34$ | $122.3 \pm 3.5$ | N/A | | $4.76 \pm 0.18$ | $124.9 \pm 3.3$ |
| $\text{K}^{+}$ | $4.91 \pm 0.24$ | $122.0 \pm 3.3$ | N/A | | $4.32 \pm 0.24$ | $128.0 \pm 3.5$ |
| $\text{Na}^{+}$ | $4.69 \pm 0.22$ | $122.9 \pm 3.2$ | $4.60 \pm 0.22$ | $124.0 \pm 3.3$ | $4.26 \pm 0.37$ | $127.7 \pm 3.6$ |
| $\text{NH}_4^{+}$ | $4.85 \pm 0.22$ | $122.3 \pm 3.3$ | $4.56 \pm 0.21$ | $124.6 \pm 3.3$ | $4.22 \pm 0.16$ | $127.8 \pm 3.9$ |

<sup>a</sup> -The conformations of the $\text{Mg}^{2+}$ -ATP complex were determined as described in the text and Fig. 4. Mean values and standard deviations of $\text{P}^{\text{A}}\text{-P}^{\text{G}}$ distance (in Å) and the $\text{P}^{\text{B}}\text{-O}^{3\text{B}}\text{-P}^{\text{G}}$ angle (in degrees) were measured over the respective parts of the simulations. Simulation periods corresponding to $\beta\gamma$ and $\alpha\beta\gamma$ conformations were identified by tracking distances between $\text{Mg}^{2+}$ and non-bridging oxygen atoms of the phosphate chain (Fig. S3); simulation periods corresponding to the "curled" conformation were identified from $\text{P}^{\text{A}}\text{-P}^{\text{G}}$ distance tracks and visual inspection of the phosphate chain shape (Fig. 3). Data for the $\alpha\beta\gamma$ coordination of the $\text{Mg}^{2+}$ -ATP complex and conformations with curled phosphate chain were calculated from simulations 1-4 in Table S3; characterization of the $\beta\gamma$ -coordination was based on simulations 5-8 in Table S3, see Table S5 for further details.

### Shape of the phosphate chain as inferred from the MD simulation data

The MD simulation data were used to compare the geometry of the ATP phosphate chain in the presence and in the absence of different M^+^ ions.

Cleavage of the bond between β- and γ-phosphates is believed to proceed via a planar transition complex, whereby the P^B^-O^3B^-P^G^ angle widens (41, 44, 51–53, 68–70). Another important feature of the Mg-ATP complex is the curvature of the phosphate chain, which can be characterized by the P^A^- P^G^ distance (Fig. 1D).

In Fig. 3, values of the P^B^-O^3B^-P^G^ angle and P^A^-P^G^ distance are plotted as a function of the simulation time.

**Figure 3.**
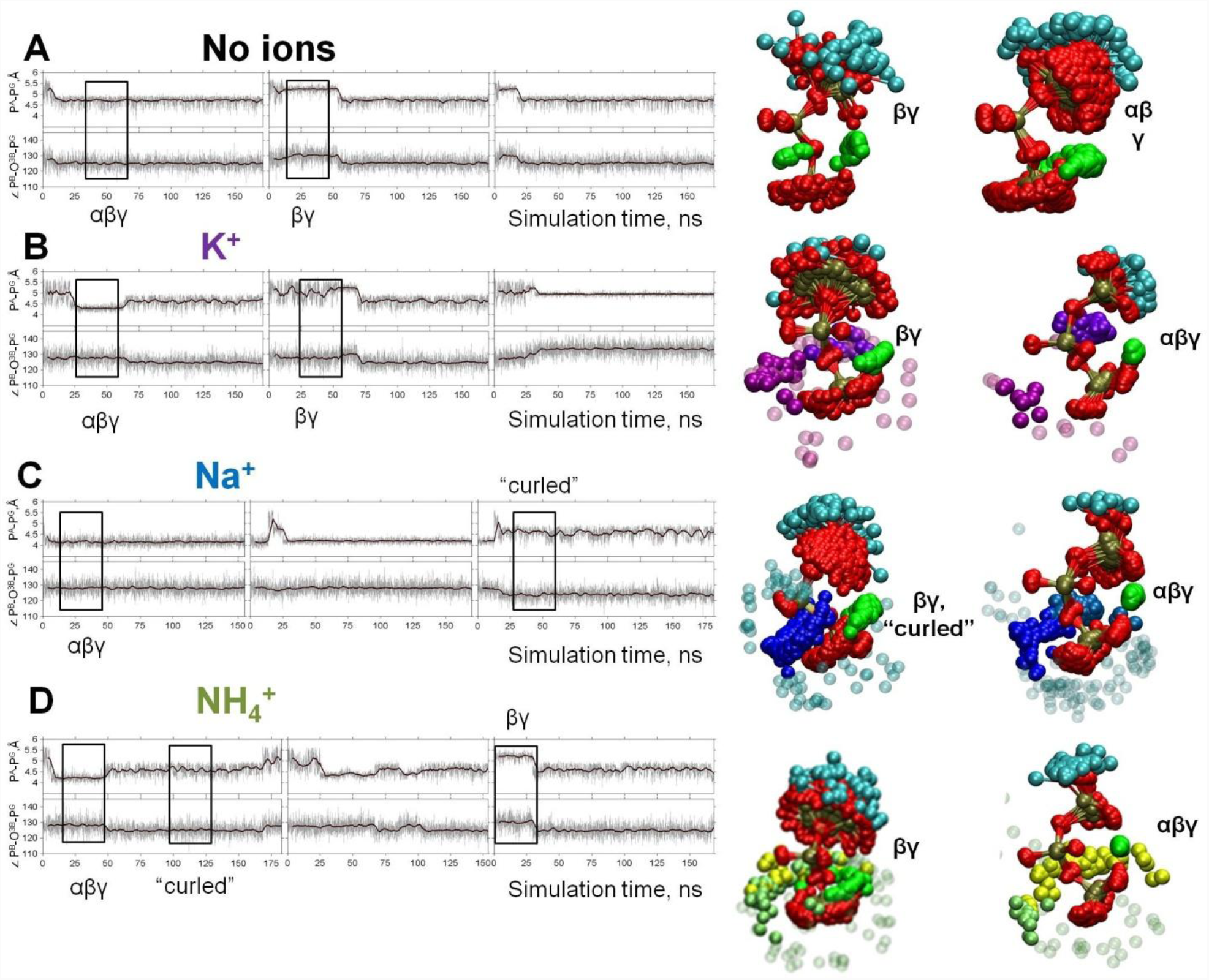
Dynamics of the phosphate chain of the Mg-ATP complex with and without monovalent cations. Each left panel shows the P^A^-P^G^ distance (upper trace) and the P^B^-O^3B^-P^G^ angle (bottom trace) in the course of MD simulations. Thin gray lines show actual values measured from each frame of the MD simulation, the bold black lines show moving average with a 2-ps window. Black boxes indicate fragments of simulations chosen for the analyses of particular types of interaction between the Mg^2+^ ion and the triphosphate chain; the respective conformations of Mg-ATP are shown on the right. The analysis was performed as in Fig. 2B. The color scheme is as in Fig. 1. A, no added ions; B–D, MD simulations in the presence of K^+^, Na^+^, and NH_4_^+^, respectively.

Fig. 4 shows heat maps of the conformations seen in the MD simulations with the values of P^B^-O^3B^- P^G^ angle and P^A^-P^G^ distance used as coordinates. The shading reflects the probability (normalized frequency) of conformations corresponding to the respective measurements.

**Figure 4.**
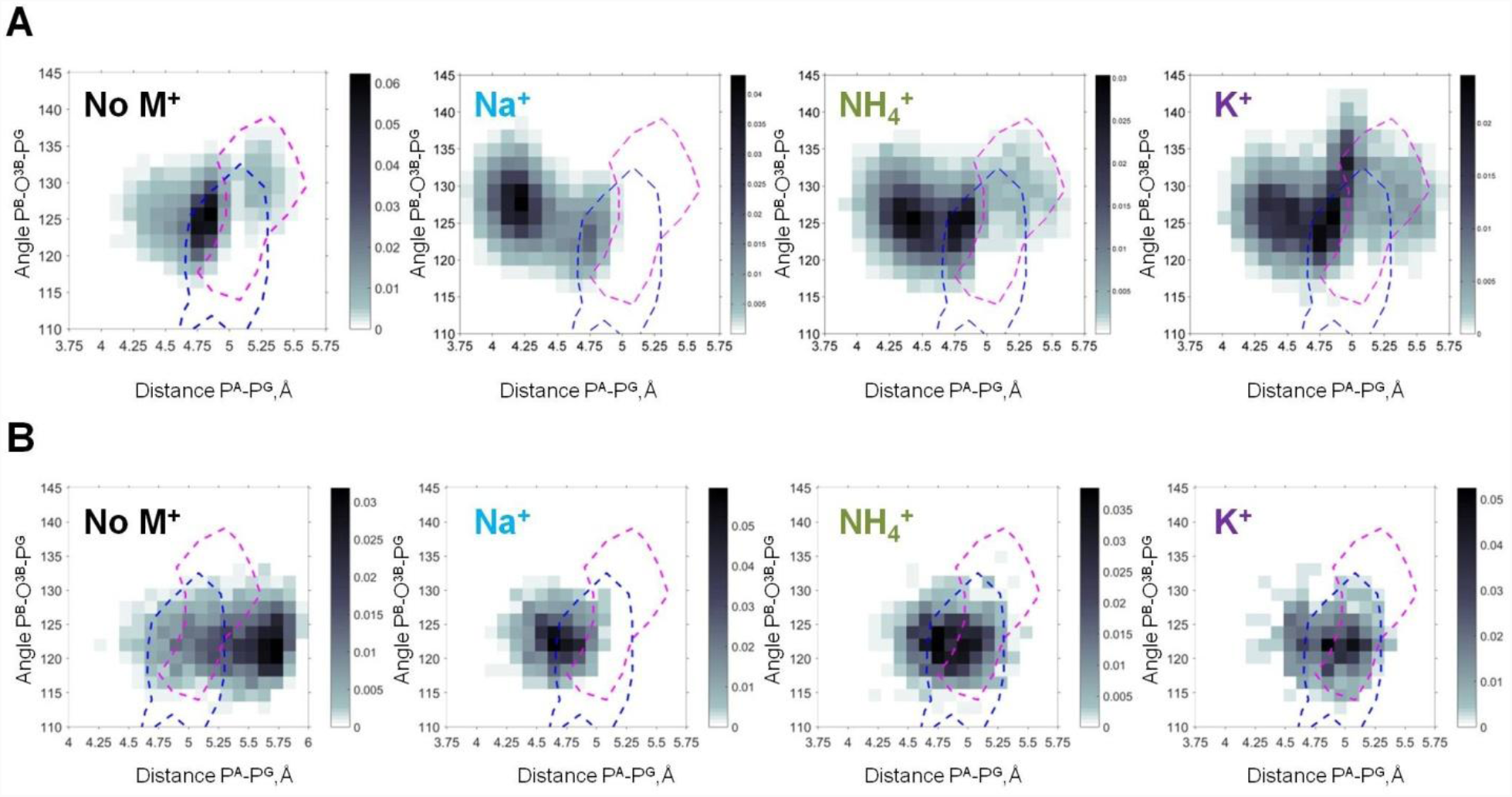
Heat maps of the Mg-ATP phosphate chain conformations distribution characterized by the P^A^-P^G^ distances (X-axis) and P^B^-O^3B^-P^G^ angles (Y-axis). Heat maps for systems with monovalent cations include only conformations of Mg-ATP complexes with at least one cation present within 4 Å radius. The color intensity is proportional to the probability (normalized frequency) of the respective conformation. Magenta dashed lines outline the areas corresponding to the conformations of transition state analogs; blue dashed lines outline the areas corresponding to the conformations of the non-hydrolyzable analogs, calculated from crystal structures of P-loop NTPases (Fig. S4). **A**. Data from the 3×170 ns simulations (no. 1 - 4 in Table S3). **B**. Data from 4×20 ns simulations of Mg-ATP in βγ conformations (no. 5-8 in Table S3, Table S5).

During all simulations, P^A^-P^G^ distances and P^B^-O^3B^-P^G^ angles fluctuated around a certain value for a while and then switched to another set of values; this behavior reflected periods of MD trajectories characterized by the same type of interaction between the Mg^2+^ ion and the triphosphate chain (Fig. 3 and S4). The ATP molecules switched between the bidentate βγ conformation and the so-called αβγ conformations with the Mg^2+^ ion being coordinated by one oxygen atom from each phosphate group (tridentate coordination of Mg^2+^). The latter conformation is known from ^31^P NMR studies (58, 71) and some proteins (72, 73). In long (3×170 ns) simulations, several versions of the αβγ conformation could be seen, differing in the particular oxygen atoms of the phosphate chain involved in the tridentate coordination of the Mg^2+^ ion (Fig. S4).

In each of the sampled conformations, the Mg-ATP complex was characterized by distinct P^A^-P^G^ distances and P^B^-O^3B^-P^G^ angles, which depended on the nature of the added monovalent cation (Fig. 3, Table 1). While all M^+^ ions seemed to contract the phosphate chain, it was more extended in the presence of K^+^ than in the presence of NH_4_^+^ or Na^+^. Furthermore, Na^+^ and NH_4_^+^ ions could induce an even more compressed, curled conformation of the Mg-ATP complex with even shorter distances between P^A^ and P^G^ atoms. Such curled conformations of the phosphate chain were not observed either in the presence of K^+^ ions or in the absence of M^+^ ions (Fig. 3, Table 1).

In short MD simulations that started from the same βγ conformation (simulations 5-8 in Table S3), we did not observe significant differences in the lifetime of the βγ conformation between systems with different cations (Table S5). For the βγ conformation of the Mg-ATP complex, the largest P^A^- P^G^ distances, up to 5.5 Å, were observed in simulations without M^+^ ions (Fig. 3, 4). Presence of M^+^ ions in the simulation system led to a significant decrease of the P^A^-P^G^ distances (Fig. 3, 4, Table 1). Among the studied cations, K^+^ ions allowed for the longest P^A^-P^G^ distances. The P^B^-O^3B^-P^G^ angles in the βγ-coordinated Mg-ATP complexes did not differ significantly between simulations with different cations or without cations added (Fig. 3, 4, Table 1).

### Shape of the phosphate chain in the structures of P-loop NTPases

Binding in the catalytic site of a P-loop NTPase imposes constraints on the Mg-NTP complex, so that only particular conformations of the phosphate chain are allowed. These conformations appear to be catalytically prone, since NTP binding to an inactive P-loop domain (in the absence of a specific activating protein) already increases the rate constant of hydrolysis by several orders of magnitude as compared to NTP hydrolysis in water (74, 75).

We analyzed the shapes of phosphate chains and the positions of positive charges around them in the available crystal structures of P-loop NTPases and compared them with the topology of Mg-ATP complexes seen in our MD simulations. The InterPro (76) entry for “P-loop containing nucleoside triphosphate hydrolase” (IPR027417) listed 2,899 X-ray and 55 solution NMR structures of P-loop proteins. From this list, we selected those X-ray structures that contain Mg^2+^ ion and an NTP-like molecule located in the proximity of at least one Lys residue, which would indicate that this NTP-like molecule is bound in the active site. Using these criteria, we identified 671 Protein Data Bank (PDB) entries, many of them with multiple subunits, resulting in the total of 1,357 Mg^2+^-NTP-like complexes. Crystal structures with non-hydrolyzable NTP analogs were used to gather information on the shape of the phosphate chain in a potentially catalytically-prone conformation. In structures with transition state analogs, the AlF_3_/AlF_4_^-^ moieties mimicked the γ-phosphate group (44, 77, 78). These structures were used as closest approximations of the nucleotide conformations in the transition state.

To characterize the conformations of the phosphate chain in the active sites of P-loop proteins, we used the same parameters as for the MD simulation data, namely the P^A^-P^G^ distance (or the corresponding distances in substrate analogs) and the value of the P^B^-O^3B^-P^G^ angle (or the corresponding angles in substrate analogs). Using these two parameters as coordinates, we mapped the conformations attained by NTP-like molecules in the crystal structures (separately shown and described in Fig. S5) on the heat maps for all four systems, calculated from MD simulations (Fig. 4). In the top row (Fig. 4A), the heat maps include all conformations of Mg-ATP in water, including those not found in crystal structures of P-loop NTPases, e.g. with αβγ coordination of Mg^2+^, as shown in Fig. 3. Therefore, conformations of Mg-ATP complexes from MD simulations only partially overlap with the conformations of non-hydrolyzable analogs of NTPs in P-loop NTPases (the blue contour in Fig. 4A). The extent of the overlap depends on the nature of the cation used in MD simulations: it is highest with K^+^ and lowest with Na^+^. The extent of this overlap was less when the data from MD simulations were compared to the conformations of transition state analogs (the pink contour in Fig. 4A). Still, in the presence of K^+^ ions, the occurrence of such transition state-like conformations was notably higher, while in simulations with Na^+^ such conformations were completely absent.

In the bottom row (Fig. 4B), we compared the conformations of the ATP phosphate chain with the βγ-coordinated Mg^2+^ ion, as obtained in the series of short (20 ns) MD simulations (Table S3) with the shapes of phosphate chains in the crystal structures of P-loop NTPases. As seen on the heat maps, in the absence of any M^+^, the phosphate chain was remarkably elongated, displaying large P^A^- P^G^ distances that were not observed either in simulations with added cations or in crystal structures. The presence of M^+^ ions led to the shortening of the P^A^-P^G^ distances. In the simulations with Na^+^ ions, the ATP phosphate chain was more contracted than in the crystal structures of P-loop NTPases (Fig. 4B). In contrast, in the MD simulations with K^+^ and NH_4_^+^ ions, the phosphate chain shape matched almost exactly the conformations of the NTP analogs in the structures of P-loop NTPases. In MD simulations in the presence of K^+^ and NH_4_^+^ ions, the distribution of the conformations of Mg-ATP complex spreads over the areas of non-hydrolyzable NTP analogs and covers even transition state analogs (Fig. 4B). Only the conformations of the transition state analogs with severely widened (>135°) P^B^-O^3B^-P^G^ angle were not matched by the MD-derived conformations.

Altogether, Fig. 4 shows that the conformational space of phosphate chain conformations, as seen in P-loop NTPases, overlapped much better with conformations seen in the MD simulations of Mg-ATP with K^+^ and NH_4_^+^ ions than with conformations obtained with Na^+^ ions.

### Cations in the active sites of P-loop NTPases

To further analyze the roles of M^+^ ions in P-loop NTPases, we selected 10 crystal structures of P-loop GTPases and ATPases (2), representing different families of P-loop proteins. We have chosen mainly the structures with non-hydrolyzable NTP analogs and transition state analogs in complex with Mg^2+^ ions, as these structures provide positions of all three phosphate groups. These structures were superposed by matching the coordinates of the P-loop regions against the structure of the K^+^- dependent GTPase MnmE [PDB: 2GJ8] (15), see Fig. 5. Each structure was then inspected to determine the locations of the positively charged residues around the phosphate chain. Fig. 5 shows that the binding sites for M^+^ ions observed in the MD simulations (Fig. 5A) were exactly those occupied by positively charged groups in the structures of P-loop NTPases (Fig. 5B, C). The binding site between the β- and γ-phosphates (the BG site) is always occupied by the amino group of the conserved P-loop lysine residue, whereas the binding site between the α- and γ-phosphates (the AG site) could be occupied, in the crystal structures, by either a K^+^ or Na^+^ ion (Fig. 5B), or an amino group of an activating lysine residue, or the guanidinium group of arginine (Fig. 5C), or a water molecule (see below).

**Figure 5.**
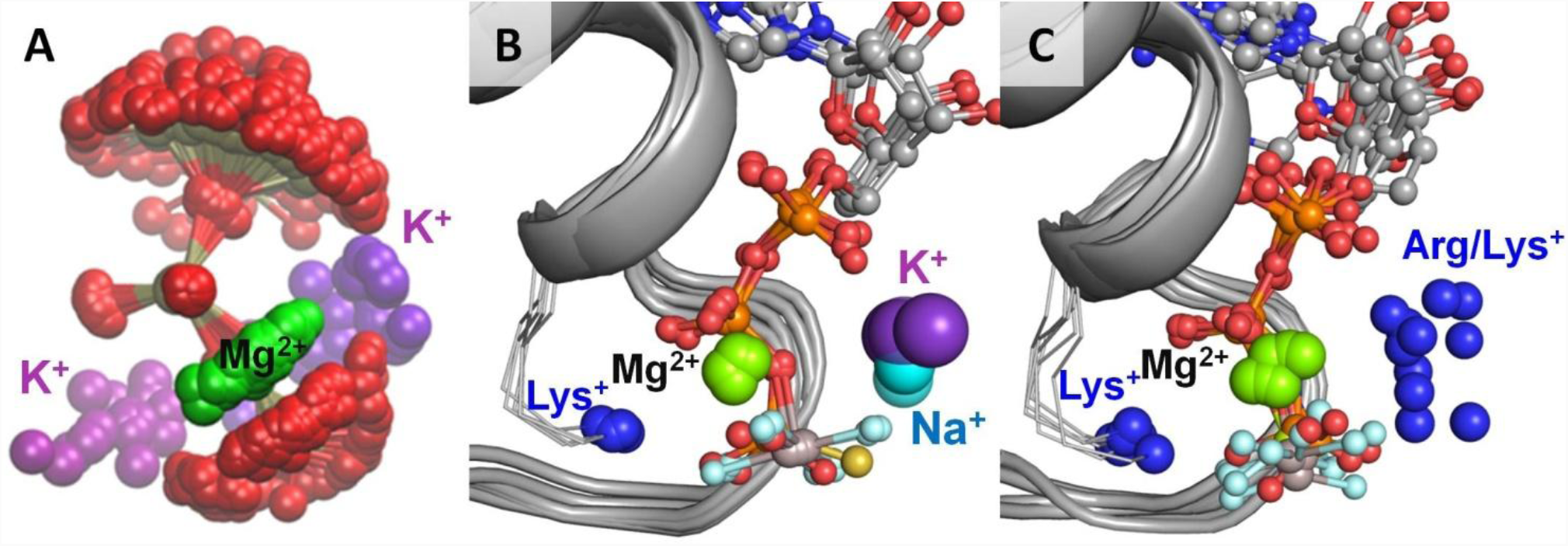
Location of positive charges around the phosphate chain of Mg-NTP complexes in solution and in protein structures. The color scheme is as in Fig. 1; blue spheres indicate positions of positively charged side-chain nitrogen atoms of Lys and Arg residues, P-loop regions are shown as cartoons in grey. **A**. Superposition of phosphate chain conformations observed in MD simulations with K^+^ ions. Only conformations with βγ coordination of Mg^2+^ are shown. **B.** Superposition of P-loop regions of crystal structures of cation-dependent P-loop NTPases: GTPase MnmE [PDB 2GJ8], Fe transporter FeoB [PDB: 3SS8], dynamin-like protein [PDB: 2X2E], and translation factor eIF-B5 [PDB 4TMZ], see Table 3 for details. **C.** Superposition of P-loop regions of crystal structures of cation-independent P-loop NTPases: Ras/RasGAP complex [PDB 1WQ1], septin [PDB 3FTQ], atlastin [PDB 4IDQ], G_α12_ protein [PDB 1ZCA], DNA polymerase III subunit τ [PDB 3GLF], F_1_- ATPase [PDB 2JDI].

In all P-loop NTPases, the phosphate chain is seen in the extended conformation similar to that observed in the presence K^+^ and NH_4_^+^ but not Na^+^ ions (Fig. 4B). Such an extended conformation is known to be stabilized by numerous interactions of all three phosphate groups with the residues of the P-loop motif, see (40) and (Shalaeva et al. submitted).

Table 2 summarizes the activation mechanisms for those classes of P-loop NTPases that contain both M^+^-activated and Arg/Lys-activated enzymes. Across different families of P-loop NTPases, different activation mechanisms have been described, usually involving interactions with other proteins, domains of the same protein, or RNA molecules, and resulting in the insertion of a positive charge - a monovalent cation or an Arg/Lys finger - into the catalytic site (13, 15–17, 24, 60, 79–81). The catalytic roles of Arg/Lys residues in the AG sites of various classes of P-loop NTPases is discussed elsewhere (Shalaeva et al. submitted). Here, we focus on the structures of P-loop NTPases that are dependent upon M^+^ ions.

**Table 2.**
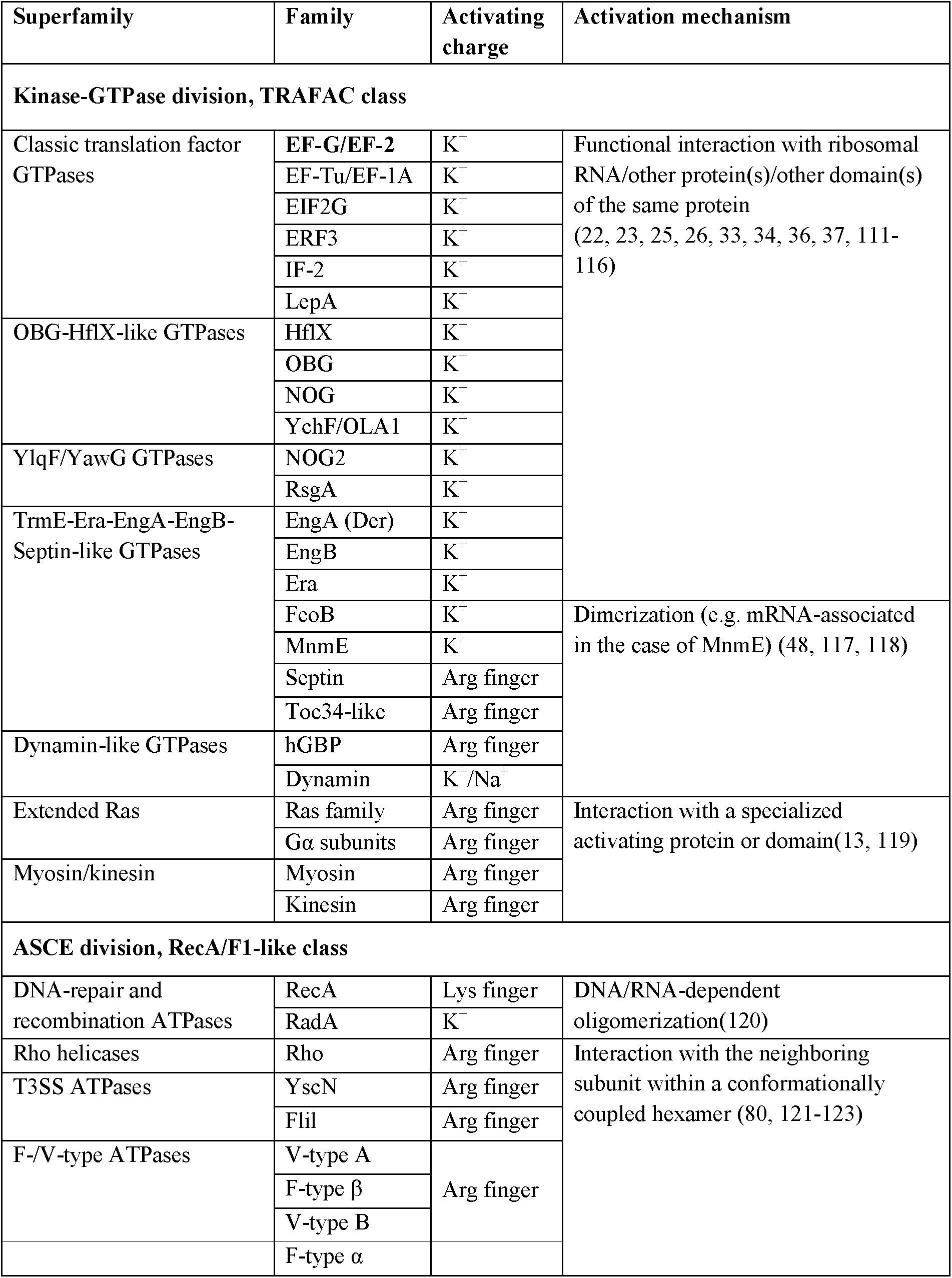
Activation mechanisms within the classes of P-loop NTPases that contain both cation-dependent and cation-independent enzymes

**Table 3.** Monovalent cation binding in crystal structures of P-loop NTPases.

| Protein | PDB<br>entry | NTP<br>analog | Charge in AG site |  |  | Phosphate chain shape |  |
| --- | --- | --- | --- | --- | --- | --- | --- |
|  |  |  | Cation | Distance<br>to P<sup>A</sup>, Å<sup>a</sup> | Distance<br>to P<sup>G</sup>, Å<sup>a</sup> | P<sup>A</sup>-P<sup>G</sup><br>distance, Å<sup>a</sup> | P<sup>B</sup>-O<sup>3B</sup>-P<sup>G</sup><br>angle, degrees<sup>a</sup> |
| TRAFAC class NTPases |  |  |  |  |  |  |  |
| GTPase | 2gj8 | GDP AlF<sub>4</sub><sup>-</sup> | K<sup>+</sup> | 2.8 | 2.6 | 5.4 | 136.3 |
| MnmE(TrmE) | 2gja | GDP AlF<sub>4</sub><sup>-</sup> | NH<sub>4</sub><sup>+</sup> | 2.9 | 2.5 | 5.4 | 136.9 |
|  | 2gj9 | GDP AlF<sub>4</sub><sup>-</sup> | Rb<sup>+</sup> | 2.9 | 2.8 | 5.5 | 131.6 |
| GTPase FeoB | 3ss8 | GDP AlF<sub>4</sub><sup>-</sup> | K<sup>+</sup> | 2.8 | 2.6 | 5.4 | 144.9 |
| Dynammin -like<br>proteins | 2x2e | GDP AlF<sub>4</sub><sup>-</sup> | Na<sup>+</sup> | 4.0 | 2.5 | 5.3 | 131.2 |
|  | 2x2f | GDP AlF<sub>4</sub><sup>-</sup> | Na<sup>+</sup> | 4.1 | 2.6 | 5.3 | 133.6 |
|  | 3w6p | GDP AlF<sub>4</sub><sup>-</sup> | Na<sup>+</sup> | 4 | 2.4 | 5.5 | 135.3 |
|  | 3t34 | GDP AlF<sub>4</sub><sup>-</sup> | Na<sup>+</sup> | 3.8 | 2.4 | 5.6 | 149.3 |
| GTPase Era | 3r9w | GNP | H<sub>2</sub>O<sup>b</sup> | 3 | 3.4 | 5.1 | 129.2 |
| Eukaryotic<br>translation<br>initiation factor<br>eIF5B | 4ncn | GTP | Na<sup>+</sup> | 2.4 | 2.4 | 5.0 | 126.6 |
|  | 4tmv | GSP | Na<sup>+</sup> | 2.4 | 2.8 (S)<sup>c</sup> | 4.9 | 126.3 |
|  | 4tmw | GTP | Na<sup>+</sup> | 2.4 | 2.4 | 4.9 | 125.9 |
|  | 4tmz | GSP | K<sup>+</sup> | 2.7 | 3.3 (S)<sup>c</sup> | 4.9 | 122.1 |
| F<sub>1</sub>-RecA-like class NTPases |  |  |  |  |  |  |  |
| DNA<br>recombinase<br>RadA | 3ew9 | ANP | K<sup>+</sup> | 6.2 | 3.3 | 5.1 | 124.5 |
|  | 2f1h | ANP | K<sup>+</sup> | 6.6 | 3.5 | 5.3 | 125.3 |
|  | 2fpm | ANP | K<sup>+</sup> | 5.9 | 2.6 | 5.1 | 124.2 |
|  | 1xu4 | ANP | K<sup>+</sup> | 6.1 | 2.7 | 5.2 | 125.0 |

<sup>a</sup> – These values were measured directly in the respective protein structures displayed in PyMOL.
<sup>b</sup> – While GTPase Era has been shown to be K<sup>+</sup>-dependent (114, 124), the crystallization solution contained no K<sup>+</sup>, only Na<sup>+</sup>, so that the likely cation-binding site is occupied by a water molecule, which forms hydrogen bonds with K<sup>+</sup> ligands.
<sup>c</sup> – Non-hydrolyzable GTP analog GDP-monothiophosphate (GSP) contains a sulfur atom in the place of the O<sup>1G</sup> atom of γ-phosphate; this atom is involved in coordination of monovalent cations in respective structures.

We have manually inspected the available structures of known K^+^-dependent P-loop NTPases (Table S1), checked for M^+^ ions bound near the NTP phosphate chain, and compared the structures of K^+^- and Na^+^-bound NTP analogs in crystal structures of P-loop proteins with the structures of the Mg^2+^-ATP-2K^+^ and Mg^2+^-ATP-2Na^+^ complexes obtained from MD simulations. In total, we were able to identify and analyze 17 structures of cation-dependent P-loop NTPases in complex with NTP analogs and K^+^, Na^+^, or NH_4_^+^ ions bound in the active site (Table 3). For each such structure, we checked the shape of the phosphate chain and the coordination sphere of the cation in the AG site. In all these structures, the distances between P^A^ and P^G^ atoms (or between the corresponding mimicking atoms) were in the range of 4.9-5.3 Å for the non-hydrolyzable analogs and 5.3-5.6 Å for the transition state analogs (Table 3). These values are similar to the P^A^-P^G^ distances observed in MD simulations of the Mg-ATP complex in the presence of K^+^ ions (Fig. 3, 4 and Table 1).

The majority of K^+^-activated NTPases, as well as the unique family of the Na^+^-adapted dynamin-related GTPases, belong to the TRAFAC class of P-loop NTPases (2), where the binding of the M^+^ ion is assisted by the so-called K-loop (20). This loop goes over the nucleotide binding site and provides two backbone carbonyl groups as additional ligands to the M^+^ coordination sphere (purple cartoon and sticks in Fig. 1B,C). To our surprise, very few structures of K^+^-dependent GTPases of the TRAFAC class contained K^+^ ions in their AG sites (*cf* Table S1 and Table 3). Furthermore, in most cases, the K^+^ loops were either unresolved or distorted (Fig. S6). Separate crystal structures with and without activating K^+^ ion were available only for the tRNA modification GTPase MnmE see Table 3 and Fig. S7. It is believed that during the catalytic turnover, two MnmE proteins undergo conformational changes to allow dimerization of their GTPase domains (G-domains) resulting in their mutual activation (16, 17). We have compared the two structures of the MnmE GTPase to further clarify their K^+^-binding determinants. In the crystallized full-length MnmE dimer, only the N-terminal domains of the two proteins interact, forming a central hinge, whereas the large helical domains and the P-loop GTPase domains (G-domains) are located on the opposite sides from the central hinge (PDB: 3GEI, Fig. S7). In such an arrangement, the distance between the active sites of the G-domains (with non-hydrolyzable GTP analogs bound) is about 20 Å (15, 16). The K-loops, responsible for cation binding, are not resolved and no K^+^ binding is observed. In the crystal structures of the isolated G-domains of MnmE in complex with the transition state analog GDP-AlF_4_^-^, which are dimerized via their K-loop (Switch I) regions (as defined in Fig. 1), the K-loops and M^+^ cations are resolved (PDB: 2GJ8, Fig. S7). The disordered K-loop in the inactive state of MnmE and the stabilized K-loop in the active state of the protein indicate that the activity of the enzyme could be controlled via formation of a full-fledged K^+^-binding site upon dimerization.

Human dynamin and the dynamin-like protein from *A. thaliana*, as noted above, are equally well activated by Na^+^ and K^+^ ions (48, 49). Dynamin-like proteins are activated upon dimerization, and crystal structures of the dimers of these GTPases in complex with GTP analogs and Na^+^ ions are available (Table 3, Fig. 1C, Fig. 6). These structures contain fully resolved K-loops, which allowed us to compare the structures of K^+^-dependent and Na^+^-adapted P-loop NTPases with the results of our MD simulations. In MD simulations, presence of Na^+^ ions led to contracted phosphate chain conformations (Fig. 3, 6A), whereas crystal structures of dynamins showed extended conformations of the phosphate chain even with a Na^+^ ion bound (Fig. 6B). In dynamin-like proteins, as in other P-loop NTPases, the phosphate chain is in the catalytically prone extended conformation owing to its stabilization by the residues of the P-loop, so that the Na^+^ ion interacts with the γ-phosphate but cannot reach the oxygen atom of the α-phosphate (Table 3, Fig. 6B). The ability of dynamins to keep the Na^+^ ion in the AG position appears to be due to the changes in the K-loop and its shortening as a result of several mutations (20), *cf* Fig. 1C and 1D. The truncated K-loop can come closer to the smaller Na^+^ ion and stabilize it in the AG position by backbone carbonyl oxygens (Fig. 6B). The non-bridging oxygen atom of α-phosphate forms an alternative hydrogen bond with the backbone amide group of Gly60 of the truncated K-loop (Fig. 6B). The truncated K-loop appears to be flexible enough to accommodate either K^+^ or Na^+^ ions, allowing dynamins to be equally well activated by K^+^ and Na^+^ ions (48, 49).

**Figure 6.**
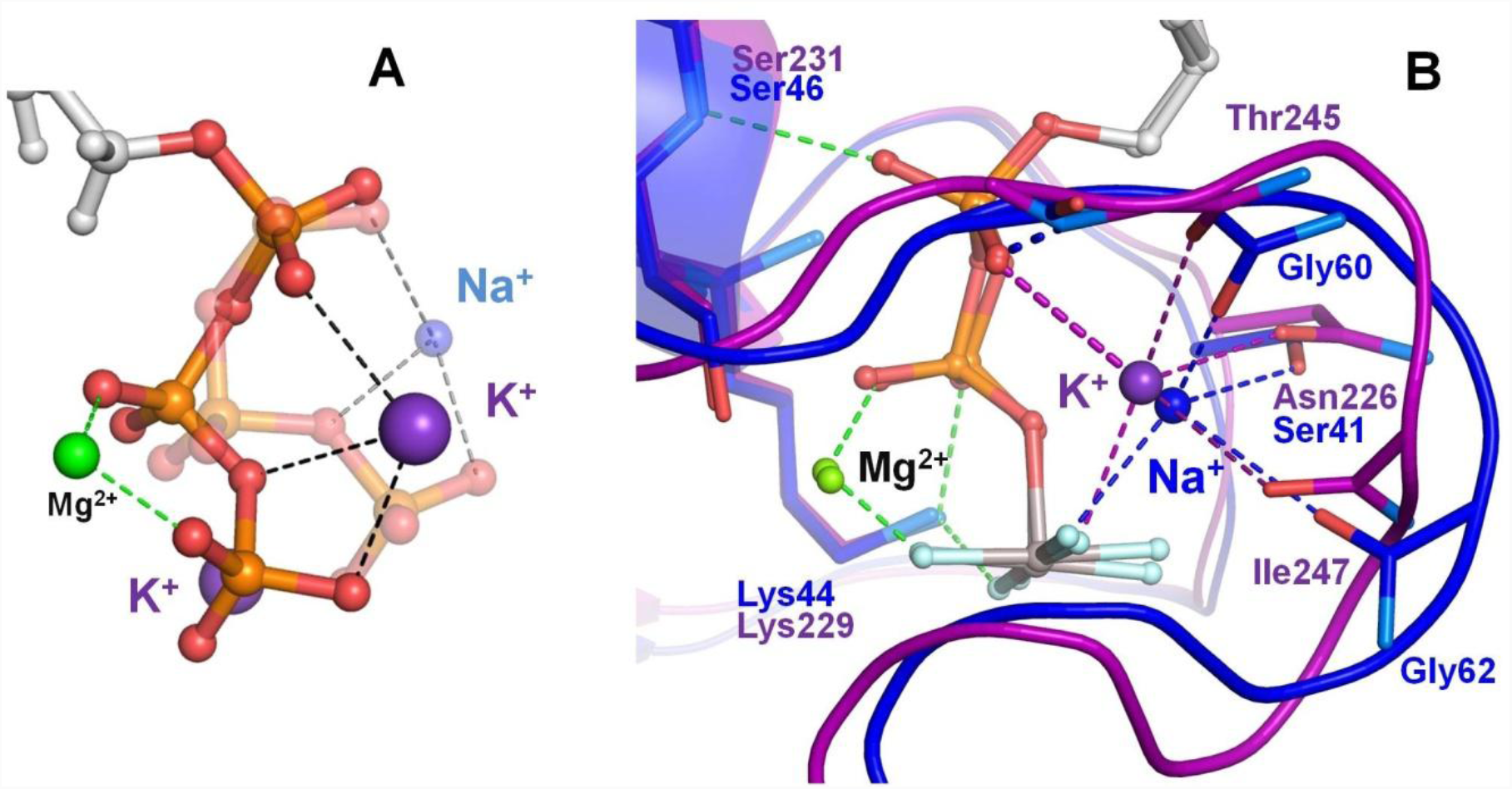
Effects of Na^+^ binding on the shape of phosphate chain in solution and in Na^+^-adapted P-loop NTPases. The color scheme is as in Fig. 1, except that Al and F atoms in the GDP-AlF_4_ complexes are colored grey and cyan, respectively. **A,** Superposition of the K^+^-bound (solid structure) and Na^+^-bound (transparent structure) conformations of the triphosphate chain as obtained from MD simulations of an ATP molecule in water. Data from MD simulations 4 - 8 in Table S3. **B**. Superposition of the P-loop NTPase structures with a bound K^+^ ion (MnmE GTPase, PDB: 2GJ8 (15), purple) and Na^+^ ion (dynamin, PDB: 2X2E (48), blue). Proteins are shown as cartoon. Dashed lines indicate hydrogen bonds and coordination bonds. Bonds that occur in all P-loop NTPases are shown in green, those that occur in K^+^-binding proteins are in purple, those bonds that occur in Na^+^- binding dynamin-like proteins are in blue. The thick dashed purple line indicates the bond between the K^+^ ion and the oxygen atom of α-phosphate, which is absent in dynamins. The thick dashed blue line indicates the dynamin-specific H-bond between O^2A^ atom and the backbone amide group of the shortened K-loop.

Outside of the TRAFAC class, only a few cases of K^+^-dependent P-loop NTPases are known, all among RecA-like recombinases (Tables 2, 3 and S1). Along with rotary ATPases, these proteins are attributed to the F_1_-RecA-like class of the ASCE (<u>A</u>dditional <u>S</u>trand, <u>C</u>atalytic <u>E</u>) division, as they bear an additional strand between the Walker A and Walker B motifs and have a conserved Glu residue in the catalytic site (2). Consequently, RecA-like recombinases are dramatically different from the TRAFAC class proteins and lack such characteristic structural motifs as Switch I/K-loop and Switch II. Crystal structure of the K^+^-dependent recombinase RadA [PDB: 3EW9] (82) shows two binding sites for K^+^ ions (Fig. S8). One of these binding sites corresponds roughly to the AG site, although the cation is shifted towards γ-phosphate and away from α-phosphate (as in the case of dynamins). The second cation is bound between γ-phosphate and the catalytic Glu residue, in the position that corresponds to the low-occupancy G-site observed in our MD simulations in water (Fig. 2A).

## Discussion

### Activation of P-loop NTPases by monovalent cations

The hydrolysis of NTPs is a key reaction in biochemistry. The high free energy of the hydrolysis is due to the repulsion between the negatively charged phosphate groups. At the same time, the cumulative negative charge of these groups repels the attacking nucleophilic groups (usually the OH^-^ ions), securing the stability of the molecule in the absence of NTP-binding enzymes (41, 83). The current views on the mechanisms of NTP hydrolysis (41, 44, 51–53, 70, 83) posit that the electrophilic γ-phosphate group is attacked by a hydroxyl group, derived from a pre-polarized water molecule. To facilitate access of the negatively charged hydroxyl to the phosphate chain, several positive charges are needed to compensate the four negative charges of the phosphate groups and, additionally, the transient negative charge of the leaving NDP group, see (41, 70) for reviews. Catalysis by known NTPases utilizes at least four positive charges: either two divalent cations (as in DNA and RNA polymerases and many nucleases and transposases (71, 84)) or a divalent cation (usually Mg^2+^) and two single positive charges in the form of (i) the conserved Lys residue and (ii) the activating M^+^ ions or Lys/Arg residues, as in P-loop NTPases. Further electrostatic compensation appears to be provided by the amide groups of the protein backbone (40), which bind and stabilize the phosphate chain in the extended conformation (Fig. 5).

The stabilization of an NTP molecule at the P-loop in an extended conformation dramatically increases the rate of hydrolysis even in the absence of an activating moiety. In Ras-like GTPases, binding of GTP to the P-loop accelerates the rate of hydrolysis by five orders of magnitude (74, 75). Delbaere and coauthors noted that, in a bound NTP molecule, the β- and γ-phosphates are in an eclipsed state owing to the interaction with the Mg^2+^ ion and conserved Lys residue of the P-loop. In this state, β- and γ-phosphates repel each other, which could explain the higher hydrolysis rate (85, 86). Hence, P-loop-bound conformations of the phosphate chains (Fig. 5) are catalytically prone.

Here, we showed that monovalent cations occupy specific well-defined sites (AG and BG sites, Fig. 2A) in the vicinity of the triphosphate chain even in the absence of enzymes. Fig. S3 shows that binding of the second K^+^ ion in the AG site can bring the phosphate chain into the fully eclipsed conformation, which has been previously suggested to be particularly catalytically productive (43). These data could explain why larger ions, such as K^+^ and Rb^+^, were shown to be more efficient than the smaller Na^+^ and Li^+^ ions in accelerating transphosphorylation even in the absence of enzymes (50), see Table S2.

The AG and BG sites are occupied by positively charged moieties in the crystal structures of P-loop NTPases (Fig. 5). The binding site between the β and γ phosphates (the BG site) is always occupied by the NH_3_^+^ group of the highly conserved P-loop lysine residue. The AG sites are usually taken by either by positively charged groups of arginine or lysine residues (Fig. 1A, 5C) or by M^+^ ions (Fig. 1B, 1C, 5B), which serve as cofactors of NTP hydrolysis (15–17, 20).

In available structures of P-loop NTPases of different classes, the phosphate chains are in the same extended conformation (Fig. 4, 5), which is maintained by the interactions with the side chains and backbone atoms of the P-loop motif, independently of the presence of monovalent activating moieties in the AG site. Thus, the P-loop keeps the phosphate chain in an extended, catalytically prone conformation, which counteracts the contracting effect of monovalent cations. In the P-loop-bound, stretched NTP molecule, a large cation, such as Arg/Lys amino group or a K^+^ ion (when stabilized by a K-loop) can coordinate both the O^2A^ and O^3G^ atoms (see Fig. 5B, S3). The smaller Na^+^-ion can neither reach both O^2A^ and O^3G^ atoms of the extended phosphate chain (Fig. 1C, 7), nor contract the phosphate chain that is fixed by the P-loop, which explains why Na^+^ ions are not competent in most P-loop NTPases.

The affinity of the AG site to K^+^ ion is intrinsically low (Table S2, Fig. 2C), therefore binding of K^+^ ions to this site in M^+^-dependent P-loop NTPases of the TRAFAC class requires a full-fledged K-loop, an extended version of the Switch I region, which provides additional ligands for the cation, see Fig. 1B, 6B and (20).

In the case of dynamins, the ability to bind either a Na^+^ ion or a K^+^ ion in the AG site was earlier traced to several mutations (20). Specifically, in dynamins, (i) the conserved Asn in the P-loop is replaced by a shorter Ser residue; (ii) the K-loop is shortened by one residue, and (iii) the Asn residue responsible for the K-loop conformation is replaced by the longer Glu residue. These mutations allow the K-loop to come closer to the small Na^+^ ion and stabilize it in the AG site even in the absence of a bond between the Na^+^ ion and O^2A^ atom (Fig. 1C, 6B). In dynamins, the free O^2A^ atom is coordinated by the backbone amide group of the shortened K-loop residue (Gly60 in the human dynamin PDB 2X2E (48)). This interaction is not seen in the structures of K^+^-dependent GTPases (*cf*. Fig. 1B with Fig. 1C). It seems that the additional coordination of O^2A^ by the Gly residue of the shortened K-loop serves as a functional replacement of its coordination by the K^+^ ion.

The observed absence of K^+^ ions from most structures of K^+^-dependent P-loop NTPases (Fig. S6) could be due to several reasons, including their absence from the crystallization medium. For example, in one of the structures of the K^+^-dependent GTPase Era, which was crystallized in the absence of K^+^ ions (PDB: 3R9W, (87)), the potential K^+^ binding site contains a water molecule (id 624) that is 2.9-3.4 Å away from six potential K^+^ ion ligands. Owing to the presence of a full-fledged K^+^-binding site, we included this structure in Table 3 (see also Fig. S9). Even when K^+^ ions were present in the crystallization medium, the electron density difference between the K^+^ ion (18 electrons) and the water molecule (10 electrons) is often insufficient to easily distinguish their relative contributions to the diffraction pattern (37). Thus, at 60% occupancy, the K^+^ ion cannot be distinguished from a water molecule (88). However, in most crystal structures of K^+^-dependent GTPases (Table S1), not only the M^+^ ion is absent, but the entire K-loop is either unresolved or shows up far away from the active site (Fig. S6). In the structures with an undefined position of the K-loop, the M^+^-binding site is incomplete, although all the sequence features of an M^+^-dependent protein, as defined by Ash et al (20), are present. Thus, other factors appear to additionally affect the K^+^ binding.

One of such factors could be inferred from the comparison of crystal structures of the cation-dependent GTPases MnmE and Era in their active and inactive conformations. A full-fledged cation binding site was absent from the inactive conformations of MnmE (Fig. S6) and Era (Fig. S9), but present in the structures where they were crystallized together with their physiological activating partners. Notably, dimerization of the G-domains of MnmE required both the GTP nucleotide and K^+^ ions in the medium, whereas Na^+^ ions could not support dimerization, even in the presence of GTP (16, 17). In the complex of Era with its activator, a 16S rRNA fragment (PDB: 3R9W), K^+^ ions were missing because of their absence from the crystallization solution. Still, the K-loop attained the shape required for cation binding and the cation-binding site was complete, with all the coordination bond partners at short distances (<3.5Å) from the water molecule that occupied the place of the K^+^ ion (Fig. S9).

The disordered K-loop in the inactive state of MnmE and Era and the stabilized K-loop in their active states suggest that the interaction with the activating partner stabilizes the functional, K^+^- binding conformation of the K-loop, which enables binding of the K^+^ ion and its subsequent interaction with the NTP molecule. Indeed, proper conformation of the K-loop (Switch I region) is crucial for the cation binding, since this loop provides two backbone oxygen atoms as ligands for the cation. We believe that the same mechanism could be involved in the activation of other K^+^- dependent NTPases (Table 2), whereby the proper conformation of the K-loop and functionally relevant K^+^ binding could be promoted by interaction with the activating protein or RNA.

In RecA-like recombinases (Fig. S8), the K^+^ ion in the AG site is coordinated by a conserved Asp residue, which is responsible for the K^+^-dependent activation (89). This residue (Asp302 in PDB: 2F1H) is provided by the adjacent monomer within the RadA homooligomer that assembles upon interaction of RecA proteins with double-stranded DNA. Thus, in RecA-like recombinases, the K^+^- binding sites differ from those in K^+^ (or Na^+^)-dependent TRAFAC NTPases, but, similarly to TRAFAC NTPases, appear to attain functionality upon the interaction with the activating partner that provides ligands for the K^+^ ion.

In conclusion, in P-loop NTPases, the activating amino groups of Arg/Lys residues or K^+^ ions occupy the AG sites similarly to the K^+^ and NH_4_^+^ ions seen in MD simulations of Mg-ATP in water. In addition, the very formation of the M^+^-binding site next to the P-loop appears to require additional interactions of the P-loop-containing domain with activating domains or proteins, as seen in MnmE and RecA, or RNA, as seen in Era (Fig. S6-S9).

### Evolutionary implications

The major classes of P-loop NTPases appear to have emerged before the divergence of bacteria and archaea (2, 4–10, 90–92). An evolutionary scenario for the origin of P-loop NTPases has been recently proposed by Lupas and colleagues, who hypothesized that the ancestor of P-loop NTPases was an NTP-binding protein incapable of fast NTP hydrolysis, but, perhaps, involved in the transport of nucleotides (4). Indeed, as already discussed (2), the main common feature of the P-loop NTPases is the eponymous motif, which was identified as an antecedent domain segment by Lupas and colleagues (5). Milner-White and coworkers argued that the very first catalytic motifs could have been short glycine-rich sequences capable of stabilizing anions (nests) (93, 94) or cations (niches) (95); such motifs can still be identified in many proteins. Specifically, the P-loop was identified as a nest for the phosphate group(s) (93, 96). We showed here that the P-loop motif specifically binds nucleotides in the same extended, catalytically-prone conformation in different families of P-loop NTPases (Fig. 4–6, S5).

The conformational space of the Mg-ATP complex, as sampled by our MD simulations (Fig. 3,4), reflects the preferred phosphate chain conformations in water and in the presence of monovalent cations. K^+^ and NH_4_^+^ ions brought Mg-ATP into extended conformations that were most similar to the catalytically-prone conformations observed in the active sites of P-loop NTPases. It is tempting to speculate that the P-loop could have been shaped in K^+^- and/or NH_4_^+^-rich, but Na^+^-poor environments, which would favor the extended conformations of unbound (free) NTPs. Indeed, the smallest ion is this study - Na^+^ - is known to exhibit the strongest binding to the phosphate chain, which has been reproduced in our MD simulations (Table S2, Fig. S2). Consequently, tightly bound Na^+^ ions would keep the phosphate chain in a contracted/curled conformation in water (Fig. 3, 4). K^+^ and NH_4_^+^ ions are larger, which results in the wider P^B^-O^3B^-P^G^ angles and longer P^A^-P^G^ distances (Table 1, Fig. 3, 4). However, binding of K^+^ and NH_4_^+^ ions to the phosphate chain is much weaker than binding of Na^+^ (Fig. 2C-E, Table S2). Thus, stretched conformation of the phosphate chain in water could be reached in the presence of K^+^ and/or NH_4_^+^ ions only if their concentrations were distinctly higher than those of Na^+^ ions.

When an NTP molecule is bound to a P-loop NTPase, the catalytically-prone extended conformation of its phosphate chain is fully determined by the interactions with the residues of the P-loop itself. An extended phosphate chain could bind (activating) K^+^/NH_4_^+^ ions or amino groups of Lys/Arg in its AG site, but is too stretched to bind Na^+^ ions. As argued by Lupas and colleagues, one of the possible mechanisms for the emergence of diverse classes of P-loop NTPases could be a combination of the same “original” NTP-binding P-loop domain with different partners that could promote the insertion of an activating moiety into the active site (4). This suggests that K^+^ ions and/or amino groups were available as activating cofactors during the emergence of P-loop NTPases. Hence, the P-loop motif itself may have been shaped by the high levels of K^+^ and/or NH_4_^+^ ions in the habitats of the first cells. Since the emergence of the P-loop motif happened at the very beginning of life, when the ion-tight membranes were unlikely to be present, the match between the shape of the P-loop and large cations of K^+^ and NH_4_^+^ is consistent with our earlier suggestions on the emergence of life in terrestrial environments rich in K^+^ and nitrogenous compounds (38, 39).

The activating Arg/Lys residues are usually provided upon interactions of the P-loop with another domain of the same protein, or an adjacent monomer in a dimer or an oligomer, or a specific activating protein, or DNA/RNA (Table 2), so that this activation can be tightly controlled, see also (Shalaeva et al. submitted). For cation-dependent TRAFAC NTPases, however, the situation is different: the cation-binding K-loop is an extended Switch I region of the same P-loop domain (Fig. 1B, 1C, 5B, 6B). If the formation of the K-loop and binding of an M^+^ ion to it were able to proceed in an uncontrolled way, then the cell stock of ATP/GTP would be promptly hydrolyzed by constantly activated M^+^-dependent NTPases. This, however, does not happen; M^+^-dependent NTPases are almost inactive *in solo* and attain the ability to hydrolyze NTPs only after binding to an activating partner. This behavior is in line with our MD simulations that indicate rather poor binding of K^+^ ions to the “naked” AG site of the ATP molecule (Fig. 2C, S1 - S4). This poor K^+^ binding manifests itself also in the need to use very high (>>100 mM) levels of potassium salts to activate the K^+^-dependent P-loop NTPases in the absence of their physiological activating proteins or RNA (33, 37). As our comparative structure analysis showed, the functional K-loop in such NTPases is distorted in the inactive (apo-) state (Fig. S6), but attains its functional shape and eventually binds the cation upon the interaction with the activating partner (Fig. S7-S9). The interaction with the activator, however, must be highly specific to prevent the activation of hydrolysis in response to an occasional binding to a non-physiological partner. It indeed seems to be specific; Table 3 lists structures of the eukaryotic translation initiation factor eIF5B in which a kind of a K-loop formed not via their functional interaction with the ribosome, but through non-physiological crystal-packing contacts (37). Although these quasi-K-loops bind different monovalent cations, the corresponding structures contain GTP molecules, indicating the absence of hydrolytic activity. In addition, the respective P^A^-P^G^ distances are shorter than those in the structures of P-loop NTPases in their active conformations (Table 3). Apparently, in addition to cation binding, some other factors may control the catalysis and prevent spurious NTP hydrolysis. Some of these factors are discussed in (Shalaeva et al., submitted).

In spite of the long evolution of P-loop-NTPases, only in a single known case, in eukaryotic dynamins, the enzyme can be activated both by K^+^ and Na^+^ ions (20, 48, 49). The adaptation to Na^+^ ions required at least 3 mutational changes in the highly conserved parts of the protein, see (20) and Fig. 6. The low probability of this combination of changes may explain why just this one case of Na^+^-adaptation is known. In contrast, Arg/Lys residues are widespread as activators of P-loop NTPases, see Table 2 and Shalaeva et al., submitted). In a few cases (e.g. in TRAFAC class NTPases) it was possible to trace how Arg residues replaced K^+^ ions in the course of evolution in different lineages (38, 39). The recruitment of an Arg/Lys residue as an activating moiety is relatively simple and makes the catalysis independent of the oscillations of K^+^ and Na^+^ levels in the cell.

### Relation to NTPases with other folds

Our MD simulations of the behavior of an unconstrained Mg-ATP complex in water showed correlations between binding of cations to the ATP molecules and their conformation. Our data provide information not only on the interaction of M^+^ ions with Mg-ATP complexes in the bidentate βγ coordination of the Mg^2+^ ion, which is typical for the P-loop NTPases, but also on their interaction with tridentate αβγ-coordinated Mg-ATP complexes (Fig. 3, 4A, S4, Table 1).

The tridentate αβγ-coordination is found, for instance, in K^+^-dependent chaperonin GroEL and related proteins. Unlike P-loop NTPases, the GroEL from *E. coli* and the related chaperonin Mm-cpn from *Methanococcus maripaludis* were inhibited by Na^+^ ions even when Na^+^ was added over K^+^ (97). In the crystal structures of GroEL, K^+^ ion was identified in the position that corresponded to the AG site of our MD simulations, *cf* the right structure in Fig. 3B with PDB: 1PQC (72) or PDB: 1KP8 (73). The P^A^-P^G^ distance for the ATP analogs is 4.4 Å in the former and 4.3 Å in the latter structure. These distances are similar to the one obtained in the MD simulations for the tridentate αβγ-coordinated Mg-ATP complexes in the presence of K^+^ ions (4.32 ± 0.24 Å); in the presence of Na^+^ ions the distance was shorter, 4.26 ± 0.37 Å (Table 1). The available structures of Mm-cpn (PDB: 3RUV and 3RUW (98)) contain only a water molecule in the AG position of the bound nucleotide; this water molecule, however, is surrounded by 5 oxygen atoms at <3Å distance, indicating the presence of a typical cation-binding site.

For GroEL, K^+^ ion was shown to increase the affinity to the nucleotide (99). It appears that the phosphate chain, unlike those tightly bound to the P-loops, retains certain flexibility in GroEL-type ATPases, so that its shape depends on the size of the monovalent cation, as it was observed in our MD simulations. Here, binding of the Na^+^ ion would lead to a contracted, supposedly, less catalytically prone conformation. Thus, Na^+^ ions added over K^+^ ions, owing to their ability to bind more tightly, would inhibit ATP hydrolysis in line with experimental observations (97). The example of GroEL-type ATPases shows that the balance between compensating the negative charge of the triphosphate chain and maintaining its catalytically-prone conformation might be important not only for P-loop NTPases, but also for other NTPase superfamilies. Accordingly, our MD simulation data may help clarify the mechanisms in other NTPases.

## Methods

### MD simulations

To investigate the effects of cation binding on the structure of the Mg-ATP complex, we have conducted free MD simulations of Mg-ATP complex in water solution alone and in the presence of K^+^, Na^+^, or NH_4_^+^ ions. Together with monovalent cations, Cl^-^ ions were added to balance the total charge of the system. For the simulation of Mg-ATP complex in water solution without additional ions, two positive charges had to be added to balance the total charge of the system. We added two dummy atoms with singe positive charges and applied positional restraints to fix the positions of these atoms in the corners of the unit cell. In all systems, the ATP position was restrained to the center of the cell by applying harmonical positional restraints to the N_1_ atom of the adenine ring.

For simulations, we used CGenFF v.2b8 parameters for ATP^4-^ and NH_4_^+^ molecules, an extension of the CHARMM force field designed for small molecules (66). We used the TIPS3P water model, which differs from other classical models in the presence of additional van der Waals parameters for interactions between water molecules (100). For the Mg^2+^ ion, we used parameters designed by Callahan et al. (101). For Na^+^ and K^+^ ions, we used parameters of Joung and Cheatham (102).

Non-bonded interactions were computed using particle mesh Ewald method with 10 Å real space cutoff for electrostatic interactions and the switching functions between 10 and 12 Å for the van der Waals interactions. The multiple time-step method was employed for the electrostatic forces; the non-bonded interaction list was constructed using a cutoff of 14 Å and updated every 20 steps. The covalent bonds involving hydrogen atoms were constrained using the SHAKE algorithm (103) (the MD integration step, 1 fs). Then the water box and ions were added; after the addition of Na^+^ or K^+^ and neutralizing ions the total ionic strength was 0.2 M.

Molecular dynamics simulations were performed in the *NPT* ensemble. Temperature was maintained at *T*= 298 K with the Berendsen thermostat using a coupling parameter of 5 ps^-1^(104). The pressure was maintained at 1 atm by the Langevin piston method with the piston mass of 100 amu and Langevin collision frequency of 500 ps^-1^(105).

After an initial 20-ns equilibration, free MD simulations were conducted for 170 ns in three independent runs (500 ns total) for each of the four systems (K^+^, Na^+^, NH_4_^+^ and no extra ions). In our calculations, we used Gromacs v.4.5.5 (106) software with MPI implementation at the supercomputer SKIF “Chebyshev” at the Computational Center, Moscow State University.

VMD (107) was used for visualization of the Mg-ATP complex conformation during the simulations. For statistical analysis of MD trajectories, we wrote a set of MatLab(108) scripts to calculate the geometrical properties of the Mg-ATP complex and to evaluate the radial distribution functions and dissociation constants of the cations bound to it.

### Protein structure analysis

For statistical analysis of the PDB structures, we used the InterPro database (76). A list of PDB IDs of P-loop proteins was extracted for the InterPro entry IPR027417 and filtered with the RCSB PDB search engine (109) to include only those structures that contained Mg^2+^ ion and one of the following molecules (in RCSB PDB chemical IDs): ATP, GTP, ANP, GNP, ACP, GCP, ASP, GSP, ADP, and GDP. We used custom MatLab scripts to measure the distances from the NTPs (or their analogs) to the surrounding Lys/Arg residues and selected only those structures with the nucleotide bound to at least one Lys (indicating that the nucleotide is indeed bound to the P-loop and the P-loop Lys residue is not mutated). Custom MatLab scripts were also used to measure the shape of the phosphate chain in each NTP-like substrate or the transition state-mimicking molecule.

To characterize M^+^-binding sites of P-loop proteins, we have searched the available literature data for cation-dependent activities of the respective proteins, with the results summarized in Table S1. For each of those proteins, we have examined the available crystal structures in order to characterize the cation binding site(s). In total, we have selected 17 structures with metal cations, ammonium ions or water molecules (Table 3). Multiple superpositions of the P-loop proteins were built in PyMOL (110) by matching coordinates of the P-loop motif together with the β-strand and α-helix flanking this loop using the PyMOL’s “super” function. Each protein was superposed to the reference structure of the MnmE GTPase structure (PDB: 2GJ8) (15). In addition to cation-dependent P-loop proteins we have chosen six cation-independent proteins from different families for comparison (Fig. 6C).

## Acknowledgments

Very useful discussions with Drs. A.V. Golovin, A. Gorfe, J. Klare, E.V. Koonin, V.P. Skulachev and H.-J. Steinhoff are greatly appreciated. We are thankful to Dr. D. Dibrova and A. Mulkidzhanyan for their help during the launching phase of this project. This study was supported by the Deutsche Forschungsgemeinschaft, Federal Ministry of Education and Research of Germany (A.Y.M.), the German Academic Exchange Service (D.N.S.), a grant from the Russian Science Foundation (14-50-00029), and the Lomonosov Moscow State University (Supercomputer Facility, Development Program). M.Y.G. is supported by the Intramural Research Program of the NIH at the National Library of Medicine.

## Supplementary Materials to Shalaeva *et al*. “Evolution of cation binding in the active sites of P-loop nucleoside triphosphatases”

**Figure S1.**
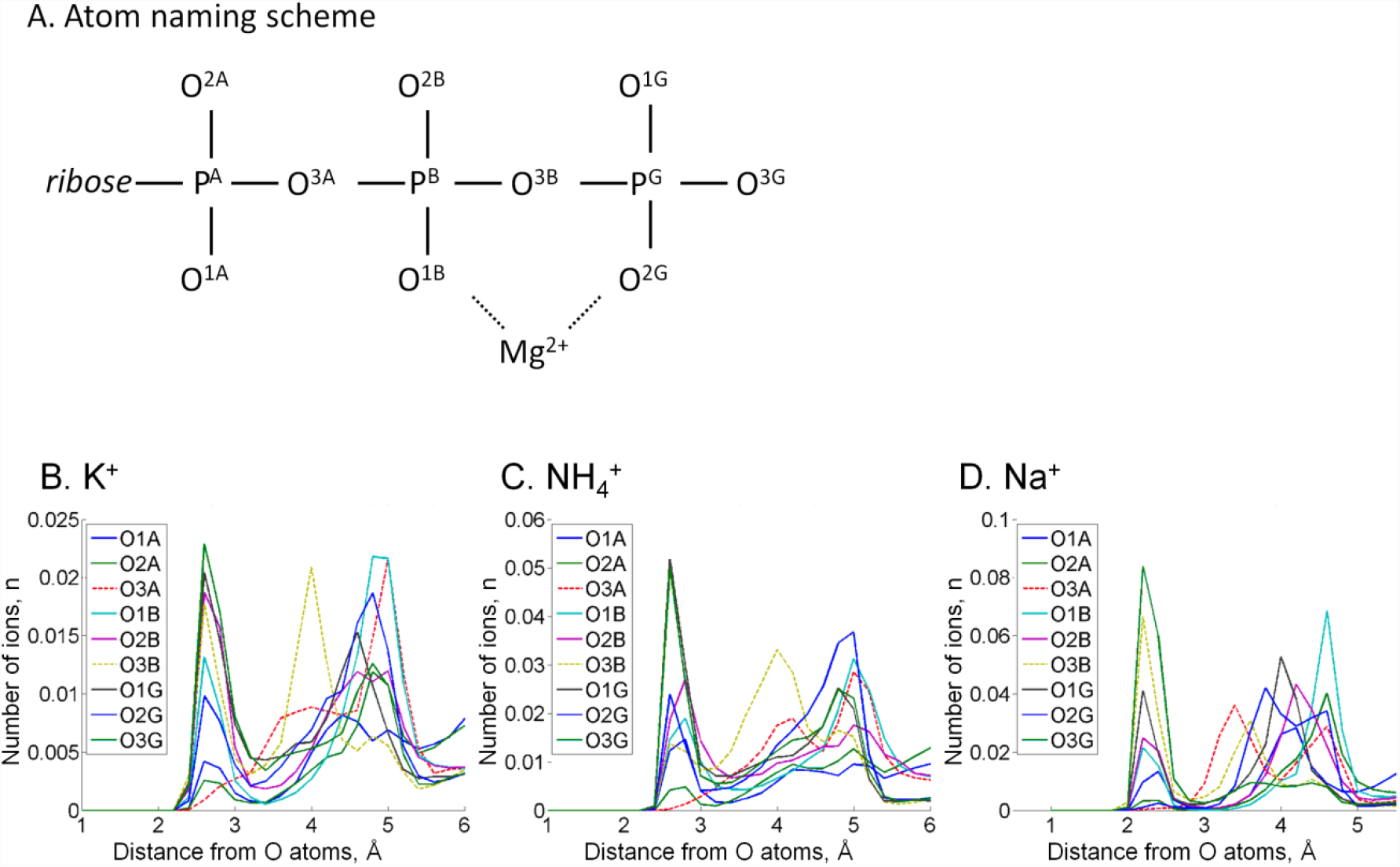
Radial distribution of cations in the proximity of each oxygen atom. Radial distributions are shown for all atoms of the ATP phosphate chain. A. Atom names according to the CHARMM naming scheme (31) and proposals in ref. (32). B. Radial distributions of cations around individual oxygen atoms. The distributions of cations around ester bond oxygen atoms O^3A^ and O^3B^ are shown by dashed lines. The peak distances from the cation to the oxygen atoms were the same 2.7 Å for K^+^ and NH_4_^+^ ions, while for Na^+^ this distance was 2.2 Å. For the NH_4_^+^ ion, the distance was measured from each oxygen atom to the nitrogen atom of NH_4_^+^. There are two ester bond oxygens in the phosphate chain, but only the oxygen (O^3B^) that connects β- and γ- phosphates was seen involved in the cation binding, it interacted more often with K^+^ and Na^+^ than with NH_4_^+^. Monovalent cations were found near oxygen atoms of γ-phosphate more often than near oxygens of β- and α-phosphates.

**Figure S2.**
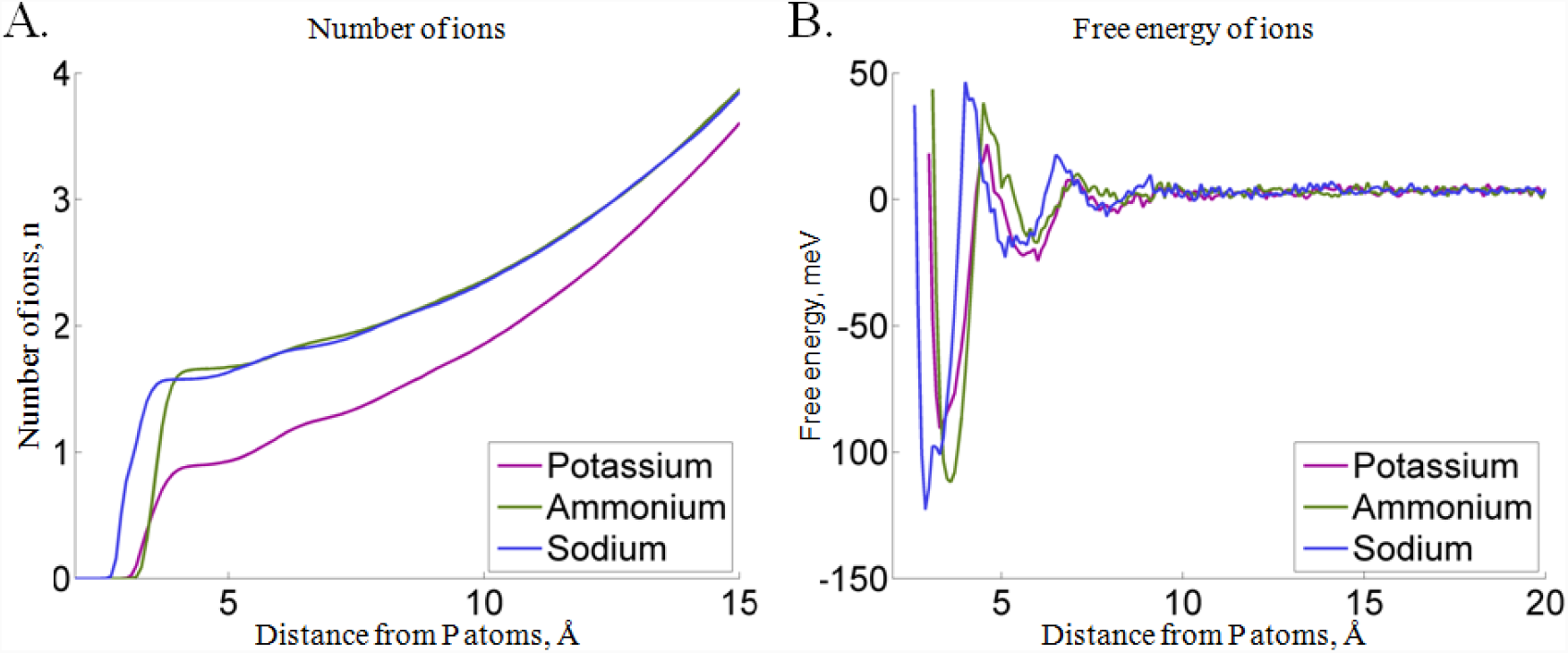
Properties of cation binding to the ATP as derived from MD simulations. (A) Probability distribution functions for cations around the phosphate chain. We have plotted the number of atoms inside the area centered on phosphorus atoms of the ATP phosphate chain as a function of the radius of the selected area. This number was estimated by measuring the distance between each cation in the system and the nearest phosphorus atom of ATP during MD simulations. The plot indicates the presence of 1.5 cations on average in the 4 Å radius around the phosphate chain in the case of Na^+^ and NH_4_^+^, and 0.75 ions on average in the case of K^+^. For all three ions, the first inflection occurs at the distances shorter than 4 Å and a less prominent second inflection can be seen at around 6 Å. (B) Free energy of the cation binding as a function of the distance from the phosphate chain, as estimated from the probability data in Fig. S2A. In addition to the two binding sites at the distances of approx. 4 Å and 6 Å, the free energy plot revealed a less pronounced third binding site at a distance of approx. 8-9 Å from the phosphorus atoms. The most prominent is the first peak, corresponding to cation binding around the phosphate chain, within the 4 Å distance of at least one of the phosphorus atoms, so further focus was specifically on cation binding around the phosphate chain.

**Figure S3.**
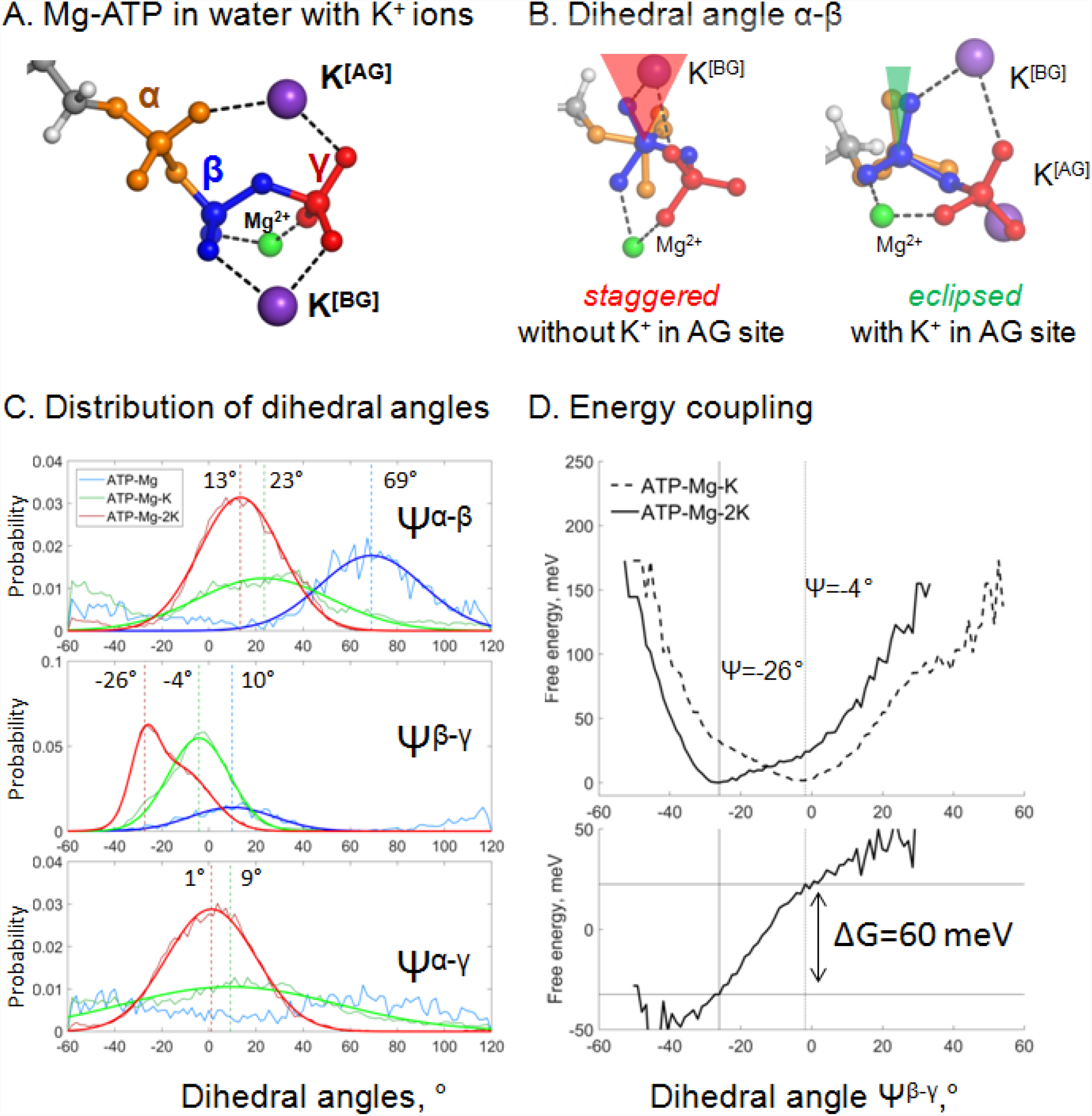
Cation binding induces eclipsed conformation of the phosphate chain. **A, B**. Mg-ATP complexes with K^+^ ions bound in the AG and BG sites; α-phosphate is shown in orange, β-phosphate in blue, γ-phosphate in red. **B.** Relative positions of α- and β-phosphates with and without K^+^ ion in the AG site; the α-phosphate is in the back, β- and γ-phosphates are in front. **C.** Distribution histograms for dihedral angles between phosphate groups in ATP, calculated from MD simulations of Mg-ATP in the absence of additional cations (blue), with one K^+^ cation bound in the BG site (green) and with two cations bound in the AG and BG sites (red). Normalized histograms of dihedral angle distribution (thin lines) were calculated from MD trajectories and fitted with normal distribution function (thick lines). Dashed lines indicate the centroid values of the fits by Gaussian function. All distributions were fitted with one-term Gaussian models, except for the Ψ^β-γ^ angle in case of Mg-ATP with two cations bound, this distribution was fitted with a two-term Gaussian, parameters for the highest peak are shown. **D.** Coupling between cation binding in the AG site and rotation of γ-phosphate relative to α- and β-phosphates. Data from 10-ns MD simulations with restraints on the positions of K^+^ ions (see the text). The top graph shows free energy calculated from normalized probabilities of ATP conformations and plotted as function of the dihedral angle between γ- and β-phosphates. The bottom plot displays free energy of coupling the binding of the second K^+^ ion with the γ- phosphate rotation, calculated as the difference between the free energy plots shown on the top graph. The lowest energy value was set to zero. These plots show that the presence of second K^+^ ion in the AG site induces a near-eclipsed state of the phosphate chain, by bringing both Ψ^α-β^ and Ψ^α-γ^ angles close to 0°, at the expense of Ψ^β-γ^, which increases slightly (see Table S4). Binding of the second K^+^ ion in the AG site stabilizes this almost eclipsed state by ∼60 meV.

**Figure S4.**
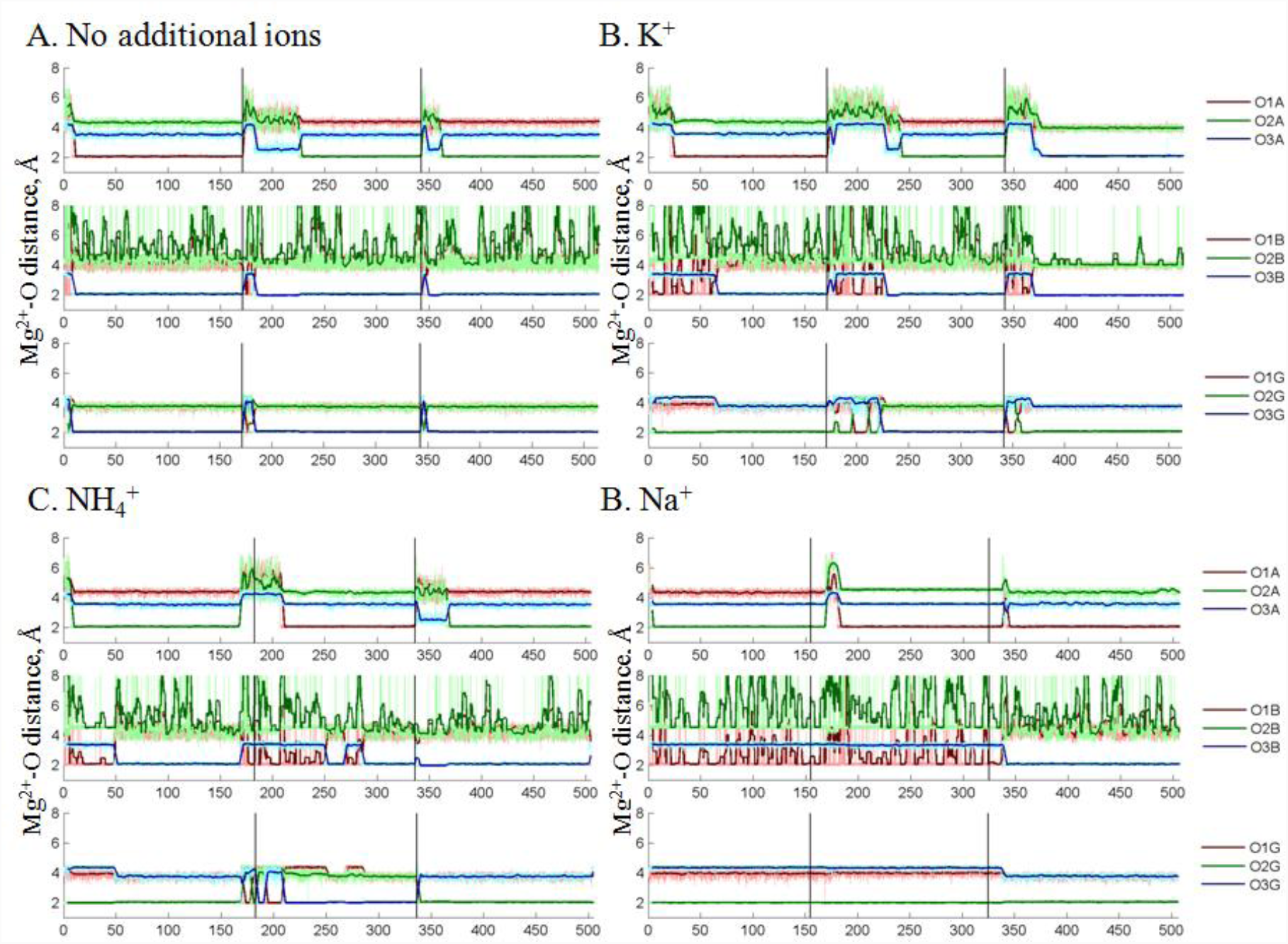
Coordination of the Mg^2+^ ion by the oxygen atoms of the ATP phosphate chain during MD simulations. Black vertical lines indicate borders between independent simulations, thick colored lines show moving average of distances measured during MD. Oxygen atoms are labelled as in Fig. S1A. The most populated conformation in each of four systems is characterized by the Mg^2+^ ion coordinated by 3 oxygen atoms: one of the free oxygens of α- phosphate (O^1A^ or O^2A^), O^3B^ atom, and an oxygen atom from γ-phosphate (O^1G^, O^2G^, or O^3G^). This conformation resembles the αβγ conformation of the Mg-ATP complex seen in other studies, but differs in the inclusion of an ester oxygen atom in the Mg^2+^ coordination sphere.

**Figure S5.**
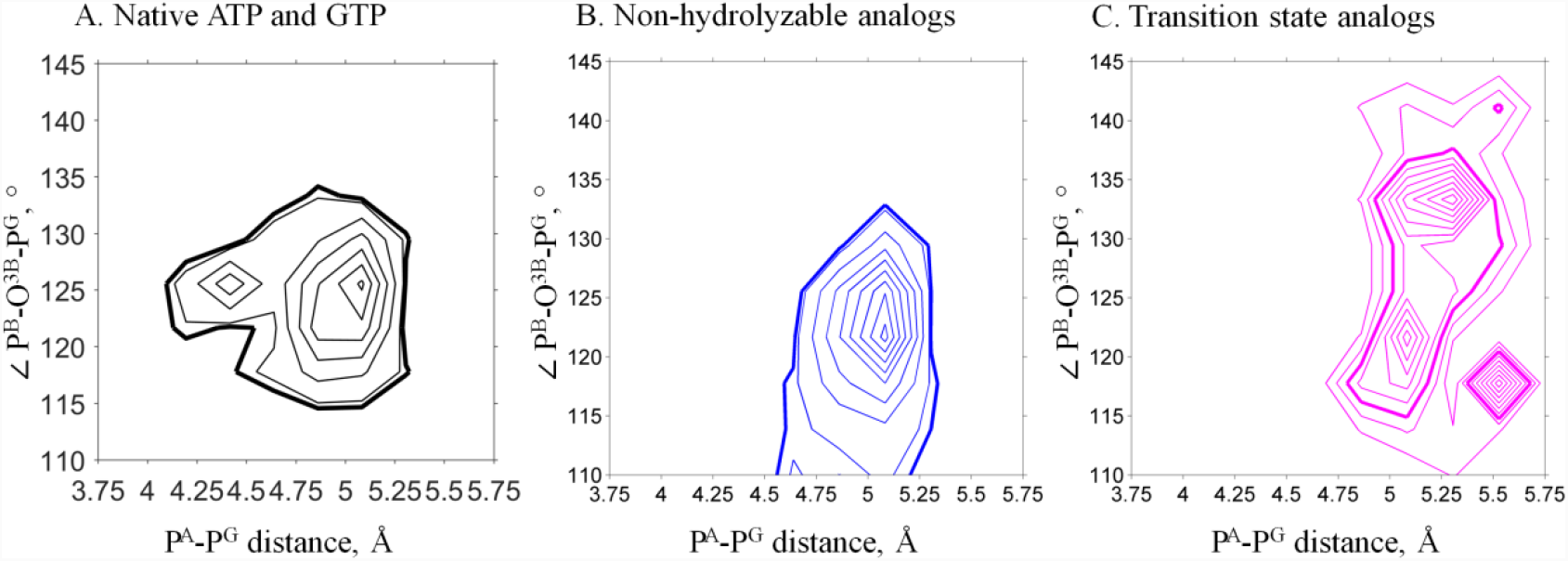
Phosphate chain shape of ATP and GTP analogs in the X-ray structures of P-loop NTPases. PDB entries for structures of P-loop NTPases were extracted from InterPro database entry IPR027417 “P-loop containing nucleoside triphosphate hydrolase” and filtered to contain only those X-ray structures that contain Mg^2+^ ions, resulting in a list of 1,333 PDB IDs. Selected structures were analyzed with custom MatLab scripts to select only those structures which contain either an NTP molecule, or its non-hydrolyzable analog, or a transition state analog. Additionally, we only considered NTP-like molecules bound in the proximity of at least one Lys residue (with less than 4.5 Å distance from NZ atom of Lys to any of the phosphate chain P atoms or the corresponding atoms in mimicking groups). In total, 1,357 NTP-like molecules from 670 PDB entries were used in the measurements. Isotherms for the heat map of the structure shape distribution are shown to indicate the most and least populated areas. Bold lines indicate isotherms chosen to represent crystallographic data in comparison with the MD results. **A.** Shapes of ATP and GTP molecules. Native ATP and GTP molecules are most likely to be crystallized with inactive proteins, so the majority of them represent non-productive conformations of the phosphate chain. **B.** Shapes of non-hydrolyzable analogs (PDB IDs: ANP, GNP, ACP. GCP, AGS, GSP). Non-hydrolyzable analogs cover lower values of the angle that is analogous to the P^B^-O^3B^-P^G^ angle, since in such molecules, the ester oxygen between P^B^ and P^G^ is replaced with another atom (N in ANP, GNP; C in ACP, GCP); or one of free oxygens of γ-phosphate is replaced with S (GSP, AGS). **C.** Shapes of the transition state analogs (ADP or GDP in complex with AlF_3_/AlF_4_^-^, VO_4_^3-^, or BeF_3_/BeF_4_^-^). ADP-BeF_x_ complexes can attain the shape of the transition state or the ground state, depending on the geometry of the active site (33). We included this complex into the group of transition state analogs with chemically similar structures. Transition state analogs represent a variety of conformations from hydrolysis-prone conformations of the substrate, via conformations that correspond to different steps of hydrolysis, to the conformations that correspond to the separated reaction products. In the latter case, the distance between P^A^ and ‘P^G^’ exceeds 5.5Å, indicating complete separation of the γ-phosphate-mimicking group from ADP/GDP; whereas the “P^B^-O^3B^-P^G^” angle decreases due to the displacement of the former ester oxygen atom that becomes the free terminal oxygen of β-phosphate. For comparison with the MD data, we only considered the major cluster of true transition state-like conformations.

**Figure S6.**
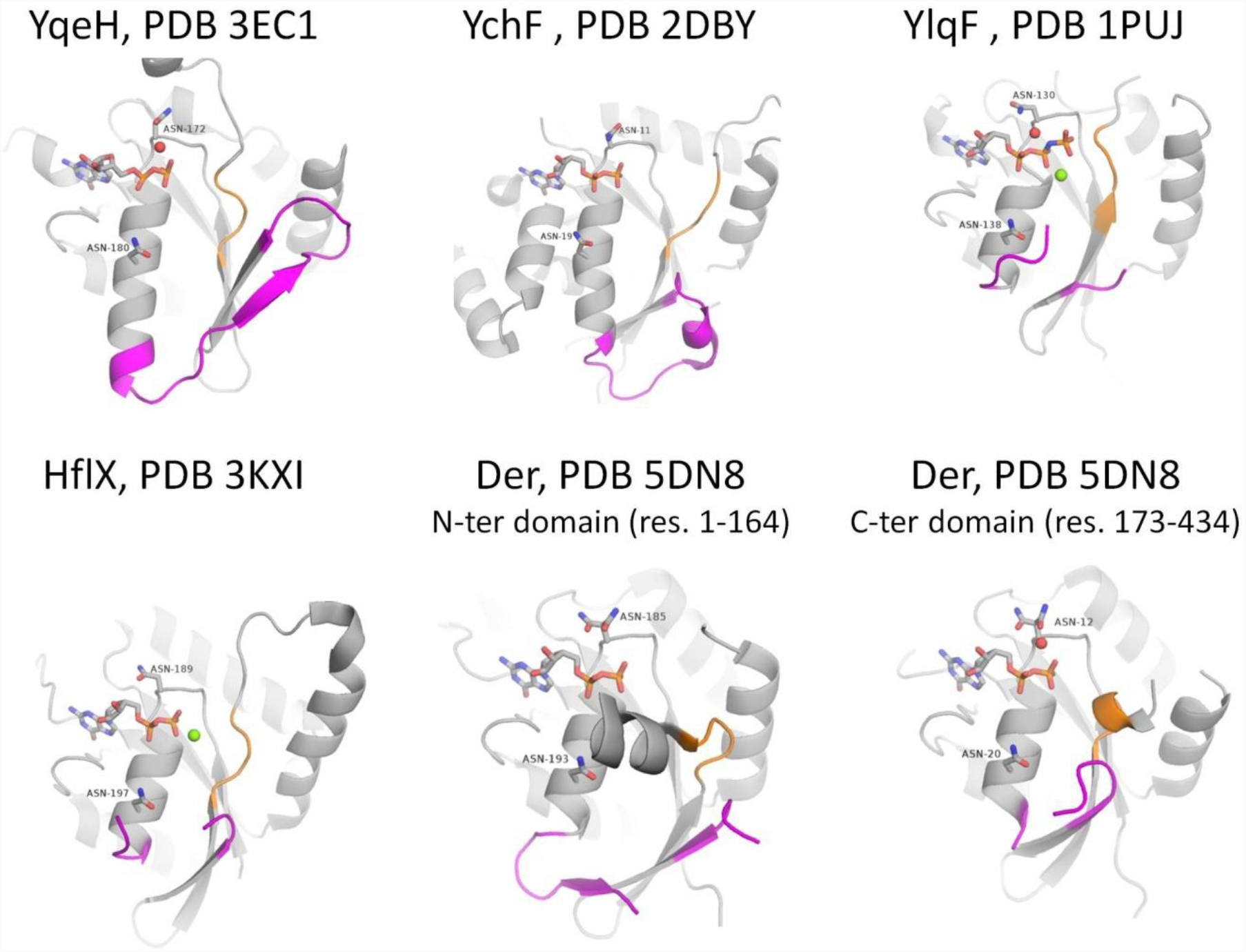
Active sites of P-loop NTPases with established K^+^-dependent activity. (see Table S1 for the full list and references). Each of the proteins shown has both Asn residues that were shown to be associated with binding of monovalent cations in related proteins (34). Switch I, including the K-loop, and its flanking regions are shown in magenta, switch II motif DxxG is shown in orange. NTP-like molecules are shown as sticks, Mg^2+^ ions are shown as green spheres, water molecules in the area of supposed cation binding are shown as red spheres.

**Figure S7.**
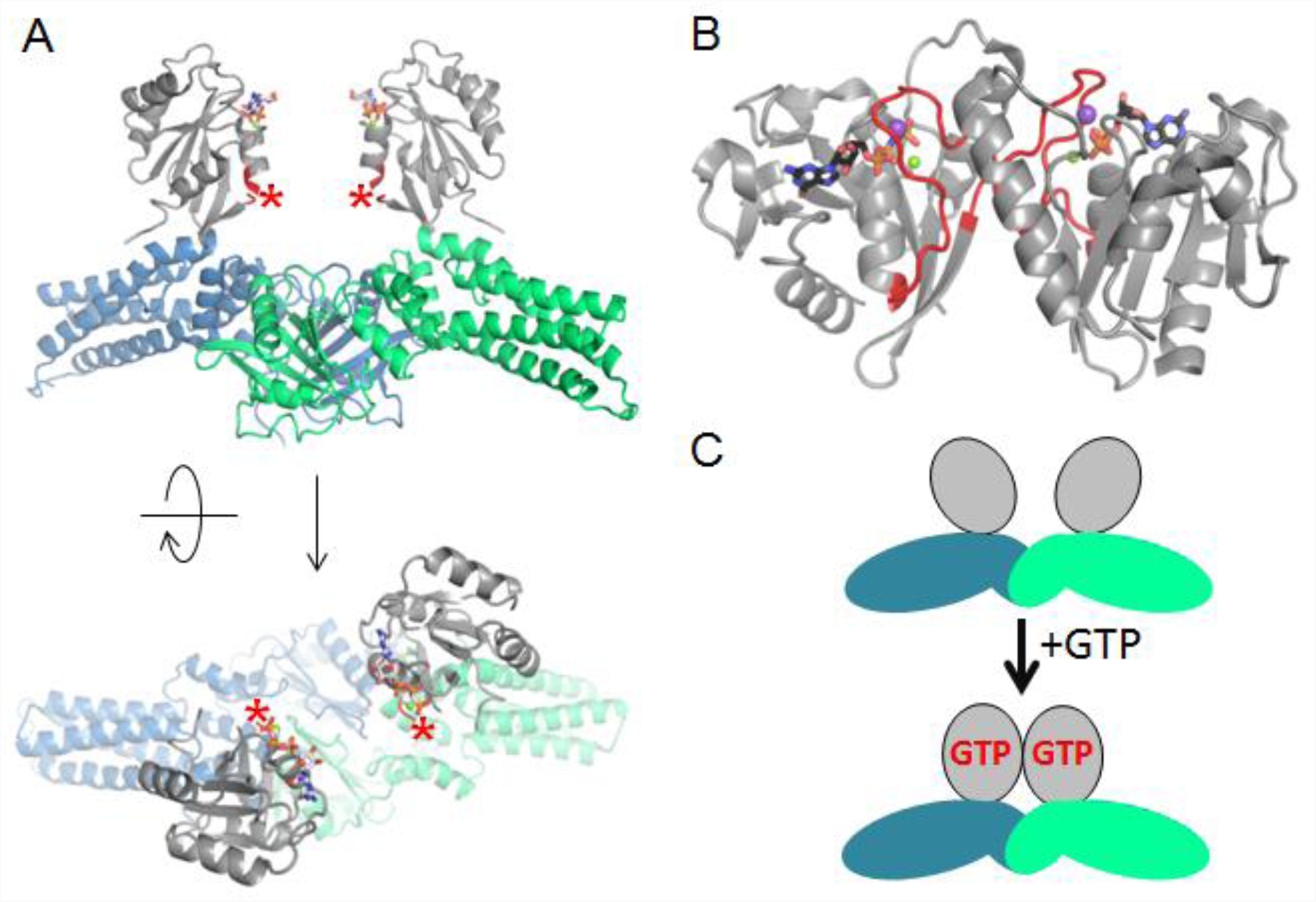
Activation of the MnmE GTPase upon dimerization. **A.** Inactive dimer of the full-length MnmE in the GTP-bound form (the structure (PDB: 3GEI) was resolved with non-hydrolyzable GTP analogs). The P-loop domain is shown in grey, the K-loop is not resolved (its position is indicated by red asterisks), the N-terminal and helical domains are shown in blue and green for different monomers. **B.** An active dimer of isolated G-domains of MnmE, as resolved in complex with a transition state analog and K^+^ ion (PDB: 2GJ8). The K-loops are shown in red, K^+^ ions are shown as purple spheres. **C.** Schematic representation of the conformational changes in MnmE dimers, reproduced after (35), domains are colored the same way as on panel A.

**Figure S8.**
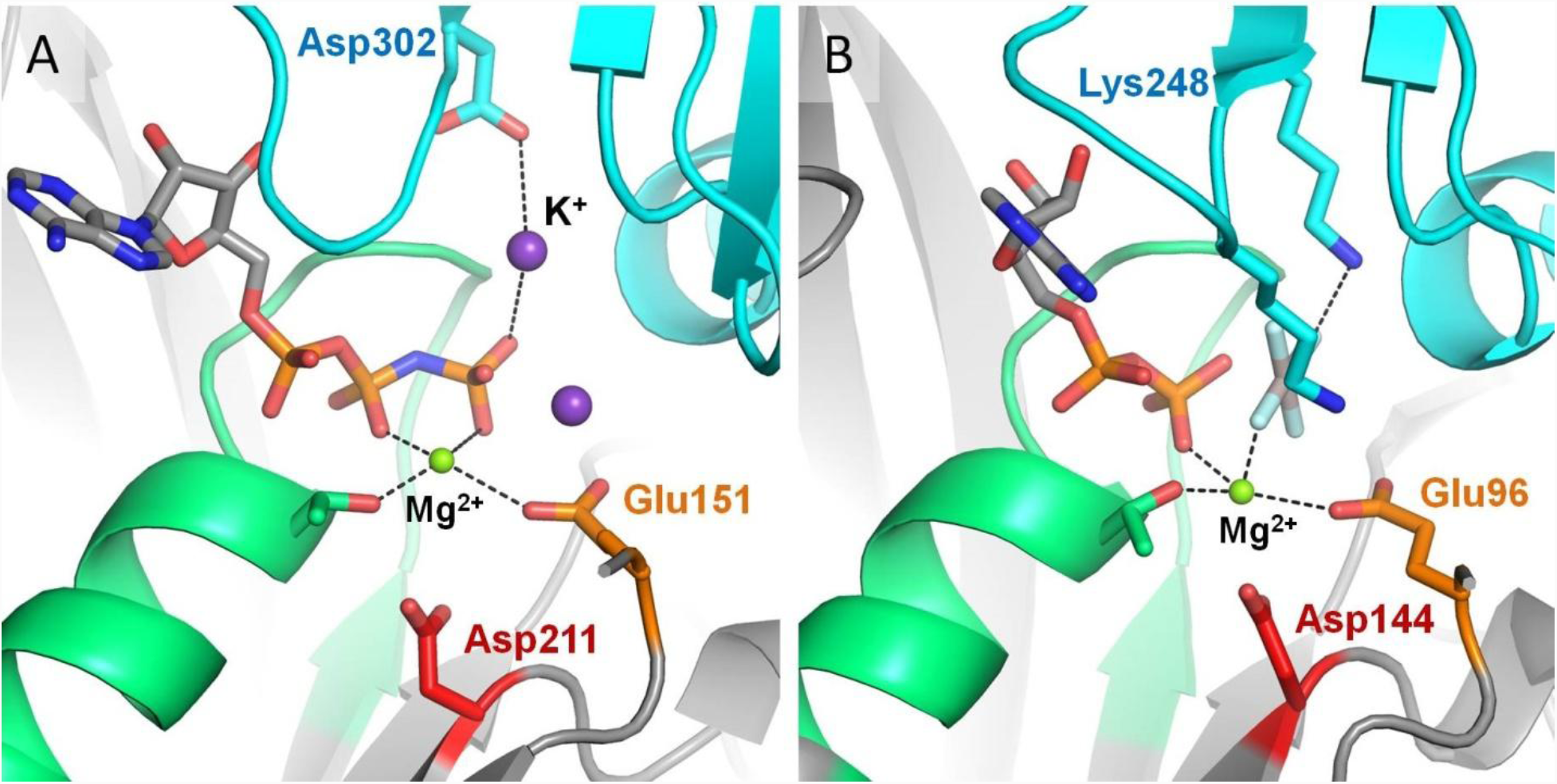
Positively charged moieties in the active site of RecA-like recombinases. **A.** Cation-dependent RadA recombinase from *Methanococcus voltae* [PDB: 2F1H] (36). **B.** Cation-independent RecA recombinase from *E. coli* [PDB: 3CMX] (37). The protein structure is shown as grey cartoon, the adjacent monomer is shown in blue, the P-loop region is shown in green; catalytic Glu residues are shown as orange sticks, conserved Asp residues of the Walker B motif are shown as red sticks. Functionally relevant residues from adjacent monomers are shown as blue sticks. Mg^2+^ ions are shown as green spheres, K^+^ ions as purple spheres.

**Figure S9.**
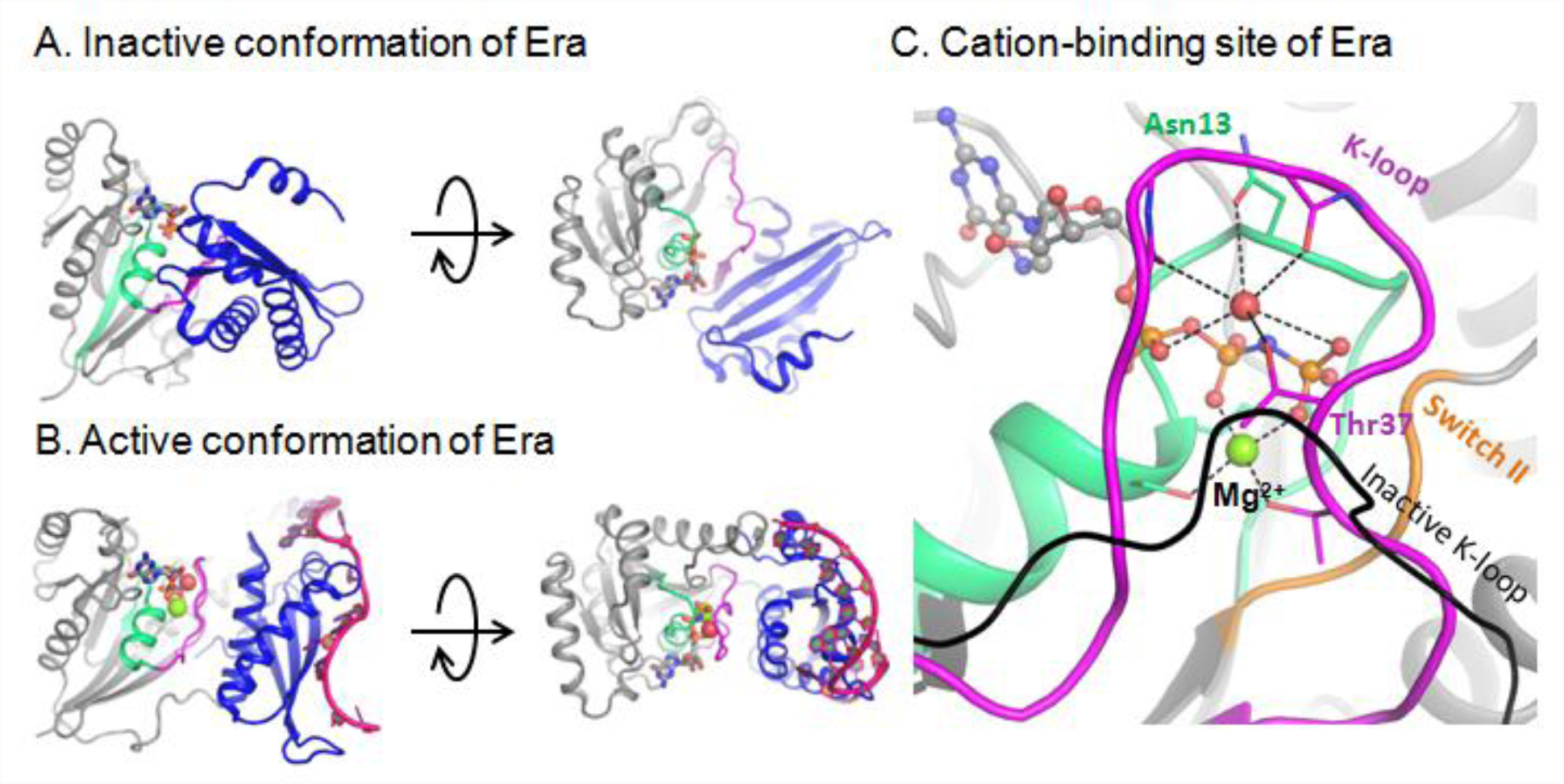
Activation of the GTPase Era upon RNA binding. **A.** Inactive Era in the GDP-bound form [PDB: 3IEU] (38) in two projections. **B.** Active Era in complex with nucleotides 1506-1542 of 16S rRNA and a non-hydrolyzable analog of GTP [PDB: 3R9W] (39) in two projections. **C.** Cation-binding site of active Era, occupied by a water molecule (shown as a red sphere) [PDB: 3R9W] (39). The black line indicates, for comparison, the position of the K-loop in the inactive structure [PDB: 3IEU] (38). The P-loop domain is shown in grey, the P-loop region shown in green, the K-loop region shown in magenta, nucleotide analogs are shown as sticks, Mg^2+^ ions are shown as green spheres.

### Statistical Analysis

To analyze conformations of Mg-ATP complexes in the presence of different cations, we selected MD simulation fragments of similar length with the same type of interaction between the Mg^2+^ ion and the triphosphate chain. In each case ∼160 ns of MD simulation were taken to characterize a particular Mg-ATP conformation; if needed, results of several independent simulations were merged to collect enough data, see Fig. S10A, B and S11A, B for examples. For the MD simulation data, we calculated autocorrelation functions (Fig. S10C and S11C). Given the correlation times obtained, independent frames were extracted to calculate characteristic values for the separate conformations of ATP. For the systems without additional monovalent cations, every N-th frame was taken for the calculation, with N defined by the correlation time. For the systems with monovalent cations, only frames in which at least 1 monovalent cation was bound to the phosphate chain were taken, with at least N frames between measurements. A monovalent cation was considered to be bound when it was within a binding distance from at least one oxygen atom of the phosphate chain, with binding distances defines a s follows: 2.4 Å for Na^+^ and 3.2Å for K^+^ and NH_4_^+^.

**Figure S10.**
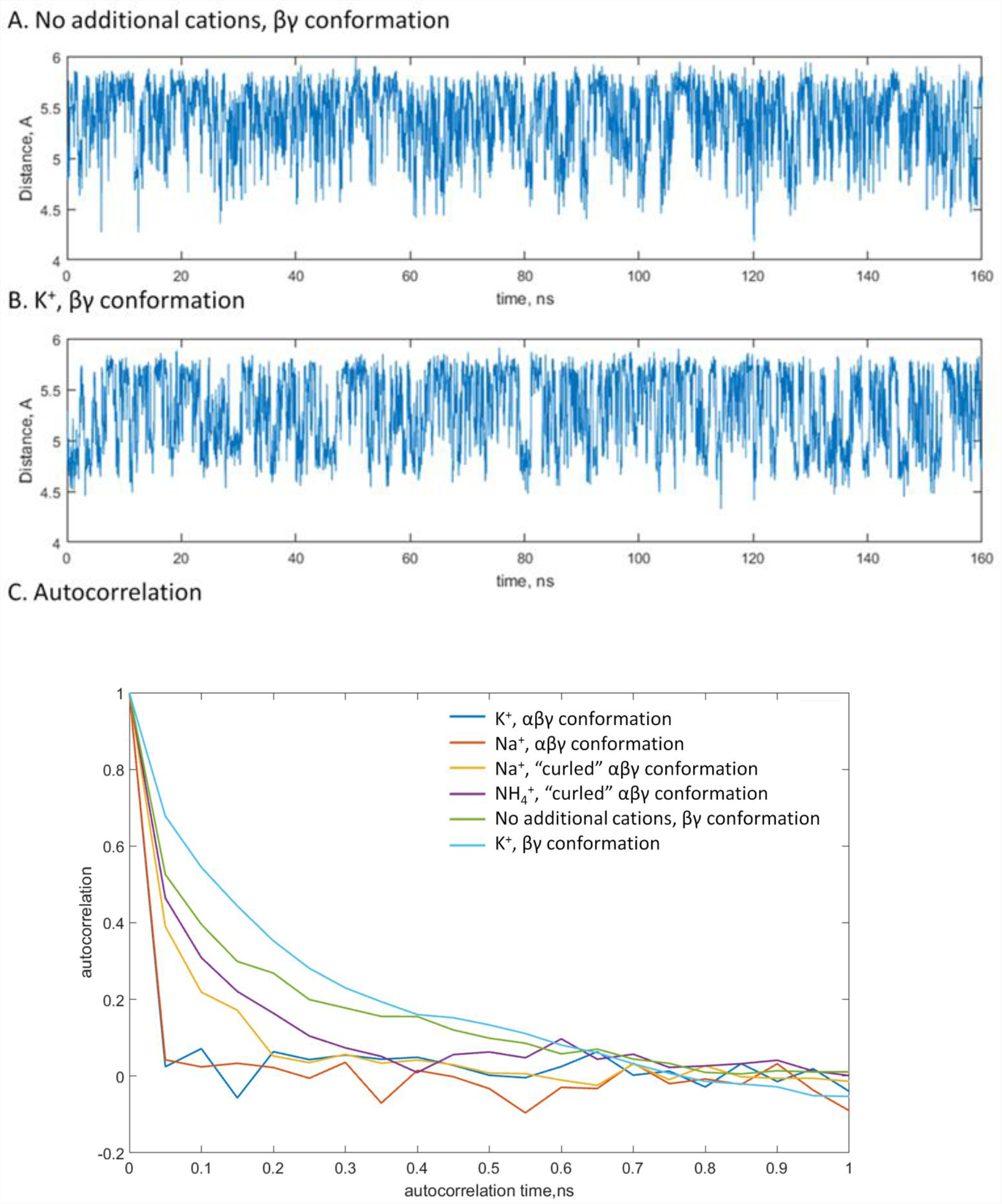
Estimation of correlation times for the Pα-Pγ distances. A,B, Changes of the distance value upon MD simulations of of βγ-coordinated Mg-ATP complexes with no additional monovalent cations (A) and with K^+^ ions (B) provided as examples; C, Autocorrelation values plotted as functions of the time lag. Based on this plot, the correlation time of 1 ns of simulation time was anticipated for the all types of interaction between the Mg^2+^ ion and the triphosphate chain and in the presence of all tested M^+^ ions.

**Figure S11.**
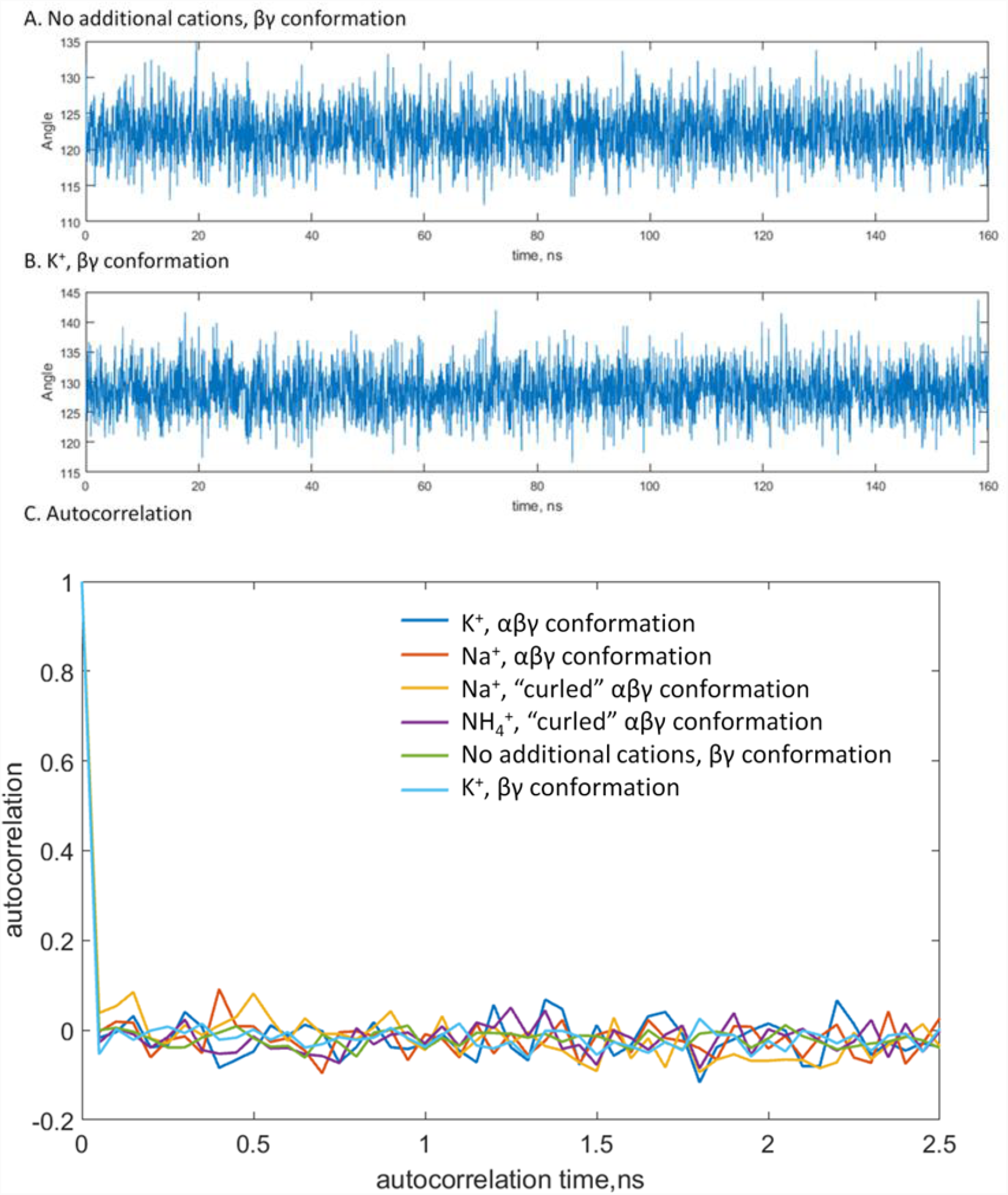
Estimation of correlation times for the Pβ-O-Pγ angles. A, B, Changes of the angle value upon MD simulations of of βγ-coordinated Mg-ATP complexes with no additional monovalent cations (A) and with K^+^ ions (B) provided as examples. C, Autocorrelation values plotted as functions of the time lag. As compared to the distance measurements, the angle values oscillated on a much shorter timescale and accordingly had shorter correlation times. From this plot, the correlation time of 5 frames or 250 ps of simulation time was estimated. The general shape of the autocorrelation function was the same for for all types of interaction between the Mg^2+^ ion and the triphosphate chain and in the presence of all tested M^+^ ions.

To compare conformations of Mg-ATP, as obtained upon MDsimulations with different monovalent cations, we used the two-sample t-test. We used the assumption that the two compared data samples were from populations with equal variances; the test statistics under the null hypothesis had Student’s t distribution with *n+m–2* degrees of freedom, where *n* and *m* were sample sizes, and the sample standard deviations were replaced by the pooled standard deviation. In each case the null hypothesis was that the data in the two samples come from independent random samples with normal distributions with equal mean values and equal but unknown variances. The alternative hypothesis was that the data in the two samples come from populations with unequal mean values. The test rejects the null hypothesis at the 5% significance level. Here we compare the test results and particular P-values obtained for all pairwise comparisons conducted in this study.

**Table S1.** Monovalent cation requirements of P-loop GTPases and ATPases

| **TRAFAC class** |  |  |  |
| --- | --- | --- | --- |
| **Protein name** | **UniProt ID** | **Cation dependence** | **Reference** |
| Dynamin-1 | DYN1_HUMAN | K<sup>+</sup>>Na<sup>+</sup> | (1) |
| Dynamin-related protein 1A | DRP1A_ARATH | K<sup>+</sup>, Na<sup>+</sup> | (2) |
| GTPase Nug1 | G0SEW3_CHATD | K<sup>+</sup>>Na<sup>+</sup> | (3) |
| Ribosome biogenesis GTPase A | RBGA_BACSU | K<sup>+</sup>, no Na<sup>+</sup> | (4) |
| Ribosome biogenesis GTPase RsgA (YjeQ) | RSGA_ECOLI | K<sup>+</sup> | (5) |
| Elongation Factor Tu, *E. coli* | EFTU1_ECOLI | K<sup>+</sup>>Na<sup>+</sup> | (6) |
| Elongation Factor Tu, *Haloarcula marismortui* | EF1A_HALMA | K<sup>+</sup>>Na<sup>+</sup> | (7) |
| Eukaryotic translation initiation factor 5B | IF2P_CHATD | Na<sup>+</sup>, K<sup>+</sup> | (8) |
| Initiation factor IF-2 | IF2_ECOLI | K<sup>+</sup> | (9) |
| tRNA modification GTPase MnmE | MNME_ECOLI | K<sup>+</sup>, no Na<sup>+</sup> | (10) |
| Ferrous iron transporter B | Q5M586_STRT2 | K<sup>+</sup>, no Na<sup>+</sup> | (11) |
| Ribosome-binding ATPase YchF | YCHF_ECOLI | K<sup>+</sup>, no Na<sup>+</sup> | (12) |
| GTPase HflX* | HFLX_BACSU | K<sup>+</sup> | (13) |
| GTPase Era | ERA_BACSU | K<sup>+</sup>, no Na<sup>+</sup> | (13) |
| GTP-binding protein EngA** *B. subtilis* | DER_BACSU | K<sup>+</sup>, no Na<sup>+</sup> | (13, 14) |
| GTP-binding protein EngA** *T. maritima* | DER_THEMA | K<sup>+</sup>, no Na<sup>+</sup> | (15) |
| NO-associated protein 1 | NOA1_ARATH | K<sup>+</sup> | (16) |
| Ribosome Assembly GTPase YqeH | YQEH_BACSU | K<sup>+</sup>, no Na<sup>+</sup> | (17) |
| Developmentally-regulated GTP-binding protein 1 | DRG1_HUMAN | K<sup>+</sup> | (18) |
| GTP-binding protein EngB | ENGB_BACSU | K<sup>+</sup>* | (13) |
| Human GTPBP3 | GTPB3_HUMAN | K<sup>+</sup> | (19) |

**Table S1. Monovalent cation requirements of P-loop GTPases and ATPases (continued)**
| **RecA-like family** |  |  |  |
| --- | --- | --- | --- |
| Human meiotic recombinase Dmc1 | DMC1_HUMAN | K<sup>+</sup> | (20) |
| Human DNA repair protein RAD51 | RAD51_HUMAN | K<sup>+</sup> | (21) |
|  |  | K<sup>+</sup>, no Na<sup>+</sup> | (22) |
| Yeast DNA repair protein RAD51 | RAD51_YEAST | K<sup>+</sup> | (23) |
| DNA repair protein RadA from *M. voltae* | RADA_METVO | K<sup>+</sup> | (24) |
| DNA repair protein RadA from *M. maripaludis* | RADA_METMI | K<sup>+</sup>, no Na<sup>+</sup> | (25) |

In the 'Cation dependence' column, 'K<sup>+</sup>' indicates that only K<sup>+</sup>-dependence has been shown; 'K<sup>+</sup>, no Na<sup>+</sup>' indicates activation by K<sup>+</sup> ions and a lack of activation by Na<sup>+</sup> ions; 'K<sup>+</sup>>Na<sup>+</sup>' denotes more effective activation by K<sup>+</sup> than by Na<sup>+</sup> ions; 'K<sup>+</sup>, Na<sup>+</sup>' and 'Na<sup>+</sup>, K<sup>+</sup>' is used when both cations have similar effects, with the more effective one listed first.
\* The GTPase activity was measured at the same concentrations of KCl and NaCl of 200 mM, and for some proteins (the second GTPase domain of EngA, HflX, EngB, all from *B. subtilis*), the lack of activation by cations has been reported (13). However, higher concentrations of ions may be required for these proteins in the absence of their activating partners, as has been shown for the second GTPase domain of EngA (14).
\*\* This protein has two P-loop GTPase domains, activity measurements were reported for the whole protein.

**Table S2.** Properties of monovalent cations and their interaction with the Mg^2+^-ATP complex.

| **Cation** | **Ionic radius (Å)<sup>b</sup>** | **Stimulation of transphosphorylation, %, <sup>a</sup>** | **Binding to ATP in the absence of Mg<sup>2+</sup> (log(K<sub>B</sub>), 25°C)** |  |  | **Binding to Mg-ATP (log(K<sub>B</sub>)<sup>f</sup>** |
| --- | --- | --- | --- | --- | --- | --- |
| Na<sup>+</sup> | 1.02 | 28 | 1.31±0.03<sup>c</sup> | 1.989±0.007<sup>d</sup> | 1.93<sup>e</sup> | **2.76** |
| K<sup>+</sup> | 1.38 | 64-73* | 1.17±0.03<sup>c</sup> | 1.873±0.005<sup>d</sup> | 1.99<sup>e</sup> | **0.88** |
| NH<sub>4</sub><sup>+</sup> | 1.44 | 27 | N/A |  |  | **1.76** |

\* measured for different salts: 64% with KCl and 73% with K<sub>2</sub>SO<sub>4</sub>.
*a* – data from (26); stimulation of transphosphorylation by 100 mM M<sup>+</sup> in the presence of 50 μM MnCl<sub>2</sub>.
*b* – data from (27)
*c* – data from (28)
*d* – data from (29)
*e* – data from (30)
*f* – calculated from MD simulations

**Table S3.** Molecular dynamics simulations performed in this work

| **No.** | **System** | **Simulation time** | **Number of repetitions** |
| --- | --- | --- | --- |
| 1 | Mg-ATP | 167 ns | 3 |
| 2 | Mg-ATP, K<sup>+</sup> | 167 ns | 3 |
| 3 | Mg-ATP, Na<sup>+</sup> | 167 ns | 3 |
| 4 | Mg-ATP, NH<sub>4</sub><sup>+</sup> | 167 ns | 3 |
| 5 | Mg-ATP | 20 ns | 25 |
| 6 | Mg-ATP, K<sup>+</sup> | 20 ns | 25 |
| 7 | Mg-ATP, Na<sup>+</sup> | 20 ns | 25 |
| 8 | Mg-ATP, NH<sub>4</sub><sup>+</sup> | 20 ns | 25 |

**Table S4.** Values of dihedral angles of the phosphate chains of Mg-ATP in the presence of K^+^ ions.

| Structure | $\Psi^{\alpha-\beta}$ | $\Psi^{\beta-\gamma}$ | $\Psi^{\alpha-\gamma}$ |
| --- | --- | --- | --- |
| Mg-ATP<br>(MD simulation) | +69±31° | +10±25° | N/A* |
| Mg-ATP-K<sup>+</sup><br>(MD simulation) | +23±40° | -4±18° | +9±65° |
| Mg-ATP-2K<sup>+</sup><br>(MD simulation) | +13±24° | -27±8° | +1±26° |

\* The rotation of α-phosphate is unrestricted and the corresponding dihedral angles can take any values between -180° and 180°.

**Table S5.**
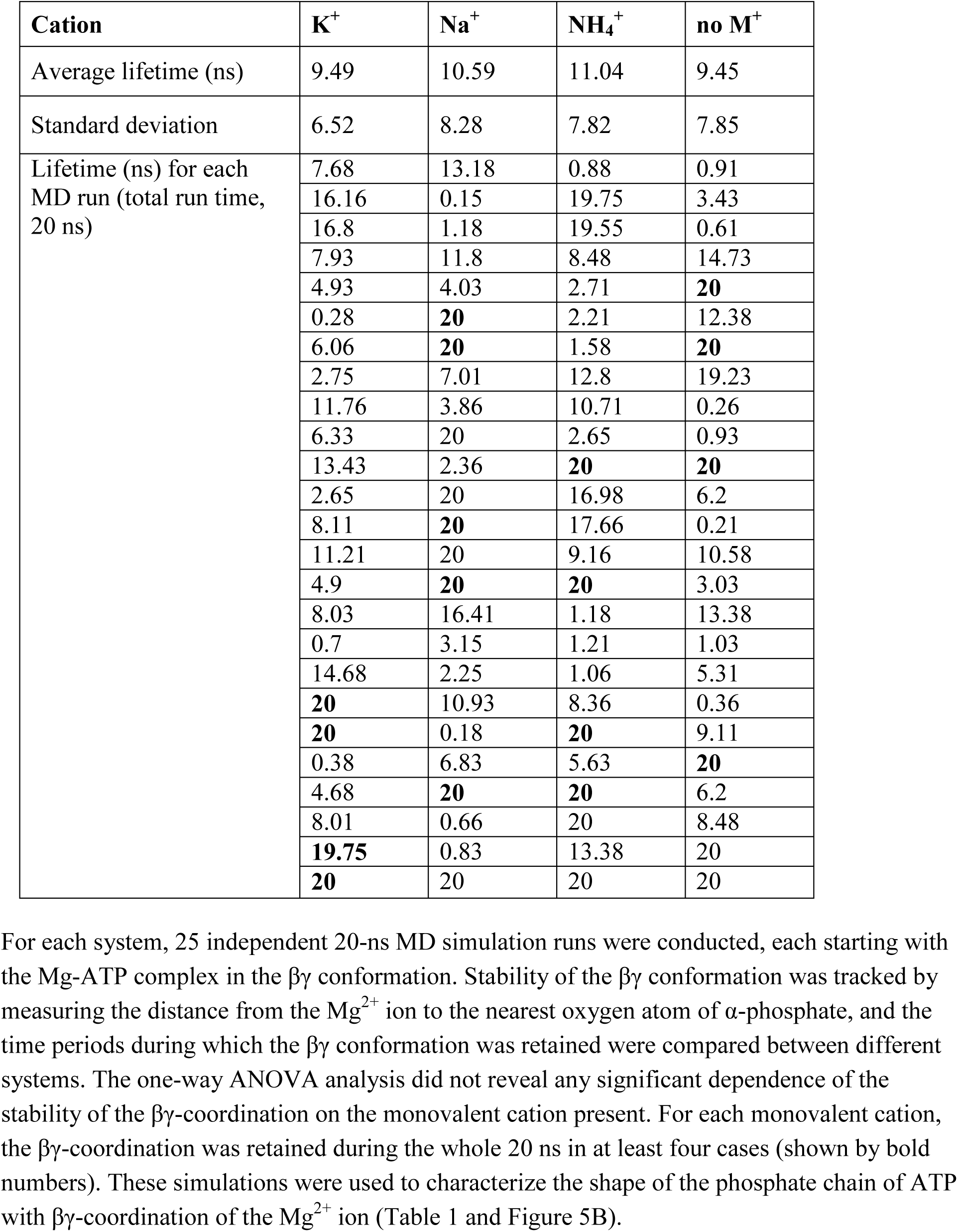
Lifetimes of the βγ-conformation of Mg^-^ATP complex during MD simulations.

| **Cation** | **K<sup>+</sup>** | **Na<sup>+</sup>** | **NH<sub>4</sub><sup>+</sup>** | **no M<sup>+</sup>** |
| --- | --- | --- | --- | --- |
| Average lifetime (ns) | 9.49 | 10.59 | 11.04 | 9.45 |
| Standard deviation | 6.52 | 8.28 | 7.82 | 7.85 |
| Lifetime (ns) for each MD run (total run time, 20 ns) | 7.68 | 13.18 | 0.88 | 0.91 |
|  | 16.16 | 0.15 | 19.75 | 3.43 |
|  | 16.8 | 1.18 | 19.55 | 0.61 |
|  | 7.93 | 11.8 | 8.48 | 14.73 |
|  | 4.93 | 4.03 | 2.71 | **20** |
|  | 0.28 | **20** | 2.21 | 12.38 |
|  | 6.06 | **20** | 1.58 | **20** |
|  | 2.75 | 7.01 | 12.8 | 19.23 |
|  | 11.76 | 3.86 | 10.71 | 0.26 |
|  | 6.33 | 20 | 2.65 | 0.93 |
|  | 13.43 | 2.36 | **20** | **20** |
|  | 2.65 | 20 | 16.98 | 6.2 |
|  | 8.11 | **20** | 17.66 | 0.21 |
|  | 11.21 | 20 | 9.16 | 10.58 |
|  | 4.9 | **20** | **20** | 3.03 |
|  | 8.03 | 16.41 | 1.18 | 13.38 |
|  | 0.7 | 3.15 | 1.21 | 1.03 |
|  | 14.68 | 2.25 | 1.06 | 5.31 |
|  | **20** | 10.93 | 8.36 | 0.36 |
|  | **20** | 0.18 | **20** | 9.11 |
|  | 0.38 | 6.83 | 5.63 | **20** |
|  | 4.68 | **20** | **20** | 6.2 |
|  | 8.01 | 0.66 | 20 | 8.48 |
|  | **19.75** | 0.83 | 13.38 | 20 |
|  | **20** | 20 | 20 | 20 |

**Table S6.** Characteristics of the triphosphate chain for different interactions between the Mg^2+^ ion and ATP.

| System | Conformation | Pα-Pγ distance, Å | Number of frames | Pβ-O-Pγ angle, ° | Number of frames |
| --- | --- | --- | --- | --- | --- |
| No cations | $\beta\gamma$ conformation | $5.4\pm0.3$ | 160 | $122.4\pm3.5$ | 640 |
| $\text{K}^+$ | $\beta\gamma$ conformation | $4.9\pm0.2$ | 85 | $128.4\pm3.5$ | 388 |
| $\text{Na}^+$ | $\beta\gamma$ conformation | $4.8\pm0.1$ | 109 | $128.2\pm3.5$ | 473 |
| $\text{NH}_4^+$ | $\beta\gamma$ conformation | $4.9\pm0.2$ | 64 | $128.6\pm3.8$ | 251 |
| No cations | $\alpha\beta\gamma$ conformation | $4.7\pm0.2$ | 161 | $125.2\pm3.3$ | 267 |
| $\text{K}^+$ | $\alpha\beta\gamma$ conformation | $4.3\pm0.1$ | 133 | $127.9\pm3.6$ | 198 |
| $\text{Na}^+$ | $\alpha\beta\gamma$ conformation | $4.2\pm0.1$ | 129 | $127.9\pm3.8$ | 192 |
| $\text{NH}_4^+$ | $\alpha\beta\gamma$ conformation | $4.2\pm0.1$ | 129 | $128.1\pm3.7$ | 190 |
| $\text{Na}^+$ | “curled” conformation | $4.6\pm0.2$ | 131 | $124.3\pm3.2$ | 194 |
| $\text{NH}_4^+$ | “curled” conformation | $4.6\pm0.2$ | 125 | $124.9\pm3.5$ | 183 |

**Table S7.**
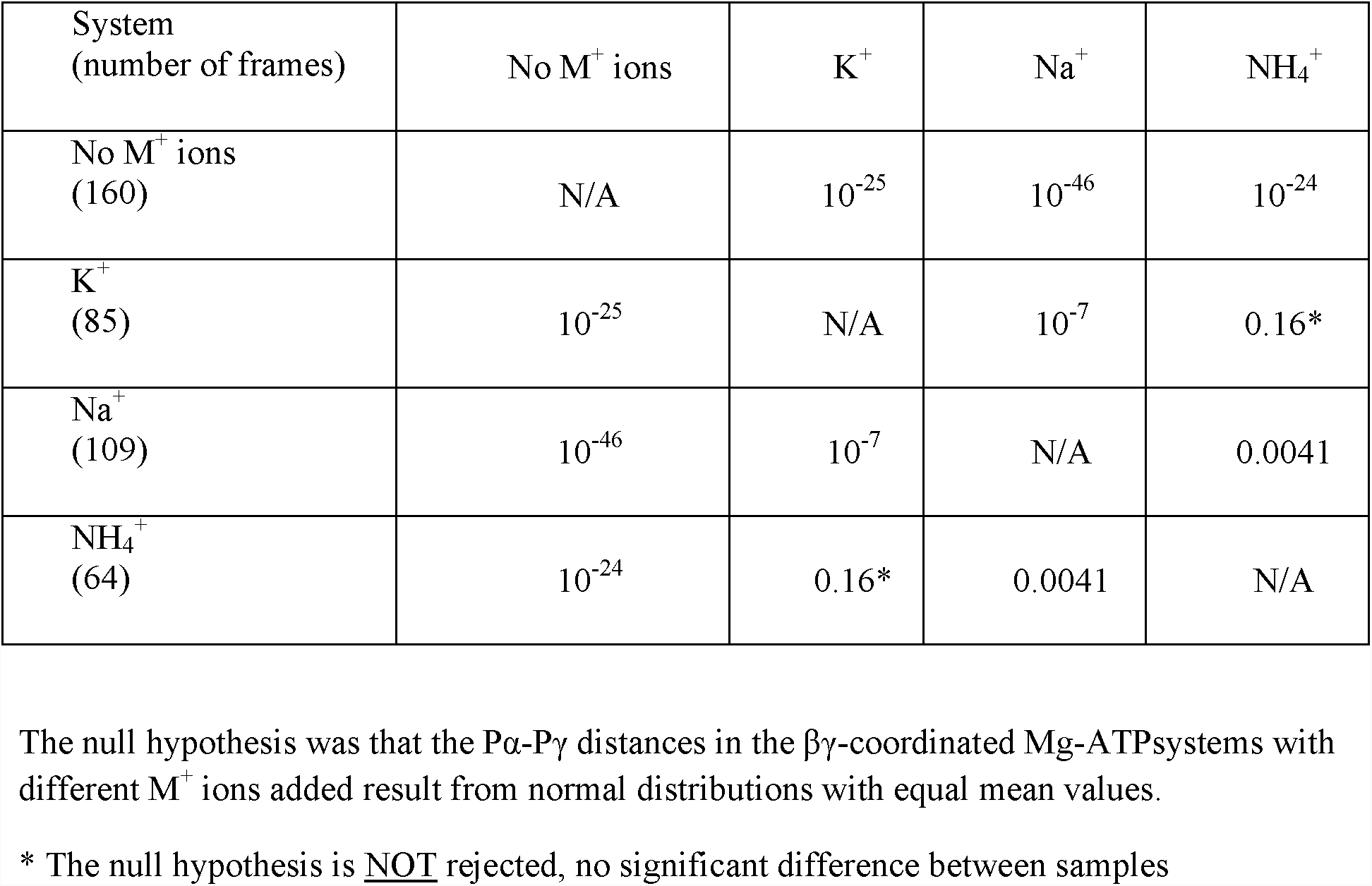
Comparison of the Pα-Pγ distance measurements of the βγ-coordinated Mg-ATP complexes

| System<br>(number of frames) | No $M^+$ ions | $K^+$ | $Na^+$ | $NH_4^+$ |
| --- | --- | --- | --- | --- |
| No $M^+$ ions<br>(160) | N/A | $10^{-25}$ | $10^{-46}$ | $10^{-24}$ |
| $K^+$<br>(85) | $10^{-25}$ | N/A | $10^{-7}$ | 0.16* |
| $Na^+$<br>(109) | $10^{-46}$ | $10^{-7}$ | N/A | 0.0041 |
| $NH_4^+$<br>(64) | $10^{-24}$ | 0.16* | 0.0041 | N/A |

The null hypothesis was that the $P\alpha$ - $P\gamma$ distances in the $\beta\gamma$ -coordinated Mg-ATP systems with different $M^+$ ions added result from normal distributions with equal mean values.
\* The null hypothesis is NOT rejected, no significant difference between samples

**Table S8.**
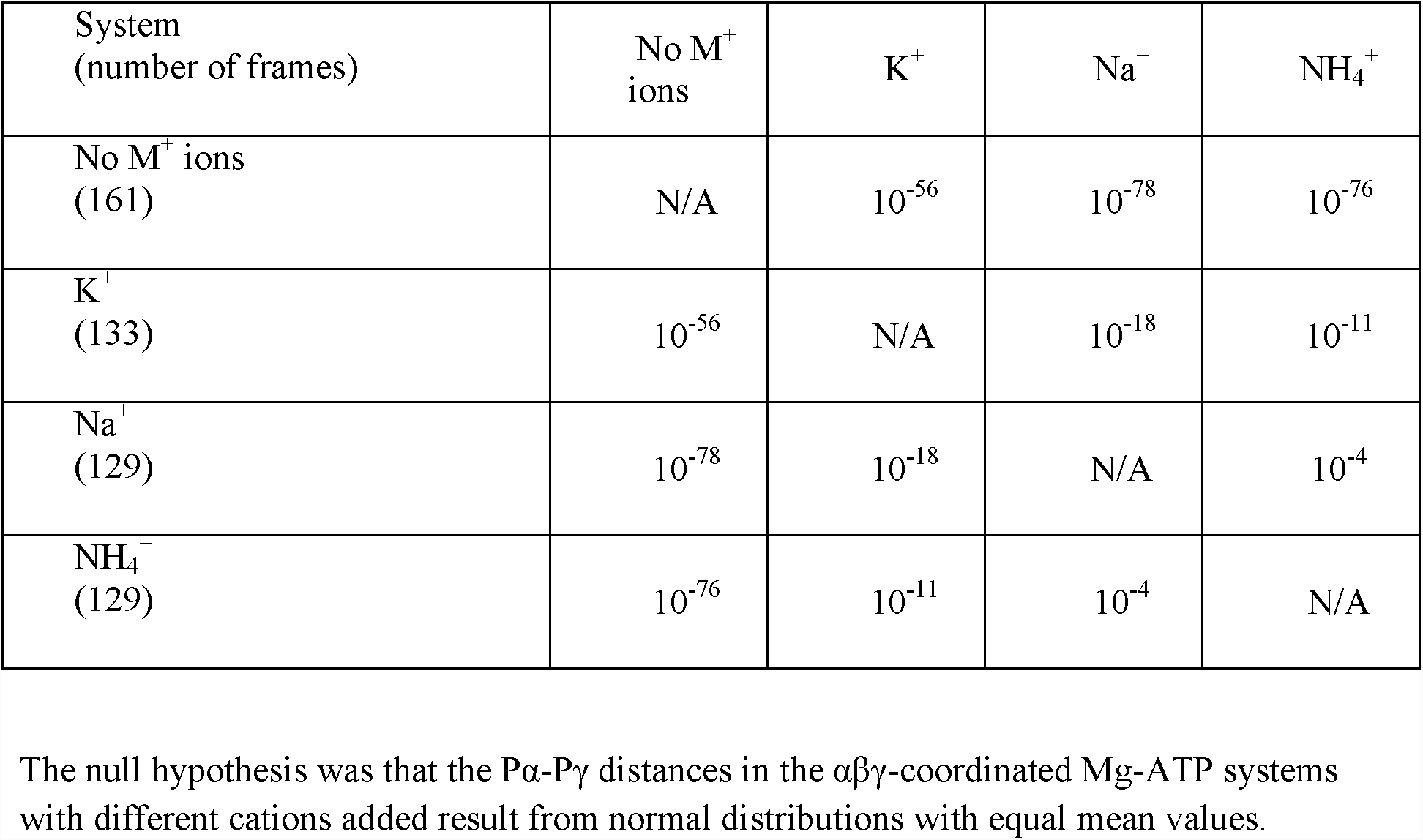
Comparison of the Pα-Pγ distance measurements of the αβγ-coordinated Mg-ATP complexes

| System<br>(number of frames) | No $M^+$<br>ions | $K^+$ | $Na^+$ | $NH_4^+$ |
| --- | --- | --- | --- | --- |
| No $M^+$ ions<br>(161) | N/A | $10^{-56}$ | $10^{-78}$ | $10^{-76}$ |
| $K^+$<br>(133) | $10^{-56}$ | N/A | $10^{-18}$ | $10^{-11}$ |
| $Na^+$<br>(129) | $10^{-78}$ | $10^{-18}$ | N/A | $10^{-4}$ |
| $NH_4^+$<br>(129) | $10^{-76}$ | $10^{-11}$ | $10^{-4}$ | N/A |

The null hypothesis was that the $P\alpha$ - $P\gamma$ distances in the $\alpha\beta\gamma$ -coordinated Mg-ATP systems with different cations added result from normal distributions with equal mean values.

**Table S9.** Comparison of the Pα-Pγ distance measurements for the αβγ-coordinated and “curled” βγ-coordinated Mg-ATP complexes in different systems

| System and conformation<br>(number of frames) | Na<sup>+</sup>,<br>$\alpha\beta\gamma$ | Na<sup>+</sup>,<br>“curled” | NH<sub>4</sub><sup>+</sup>,<br>$\alpha\beta\gamma$ | NH<sub>4</sub><sup>+</sup>,<br>“curled” |
| --- | --- | --- | --- | --- |
| Na<sup>+</sup>, $\alpha\beta\gamma$<br>(129) | N/A | 10<sup>-46</sup> | 10<sup>-4</sup> | N/A |
| Na<sup>+</sup>, “curled”<br>(121) | 10<sup>-46</sup> | N/A | N/A | 0.98* |
| NH<sub>4</sub><sup>+</sup>, $\alpha\beta\gamma$<br>(129) | 10<sup>-4</sup> | N/A | N/A | 10<sup>-42</sup> |
| NH<sub>4</sub><sup>+</sup>, “curled”<br>(135) | N/A | 0.98* | 10<sup>-42</sup> | N/A |

The null hypothesis was that the Pα -P $\gamma$ distances in ATP are the same in the $\alpha\beta\gamma$ -coordinated and “curled” $\beta\gamma$ -coordinated Mg-ATP complexes, respectively.
\* The null hypothesis is NOT rejected, no significant difference between samples

**Table S10.** Comparison of the Pβ-O-Pγ angle measurements for the βγ-coordinated Mg-ATP complexes

| System<br>(number of frames) | No M<sup>+</sup><br>ions | K<sup>+</sup> | Na<sup>+</sup> | NH<sub>4</sub><sup>+</sup> |
| --- | --- | --- | --- | --- |
| No M<sup>+</sup> ions<br>(640) | N/A | 10<sup>-118</sup> | 10<sup>-127</sup> | 10<sup>-94</sup> |
| K<sup>+</sup><br>(388) | 10<sup>-118</sup> | N/A | 0.46* | 0.49* |
| Na<sup>+</sup><br>(473) | 10<sup>-127</sup> | 0.46* | N/A | 0.17* |
| NH<sub>4</sub><sup>+</sup><br>(251) | 10<sup>-94</sup> | 0.49* | 0.17* | N/A |

The null hypothesis was that the Pβ-O-P $\gamma$ angles in the $\beta\gamma$ -coordinated Mg-ATP complexes with different M<sup>+</sup> ions added result from normal distributions with equal mean values.
\* The null hypothesis was NOT rejected, no significant difference between samples

**Table S11.** Comparison of the Pβ-O-Pγ angle measurements for the αβγ-coordinated MgATP complexes

| System<br>(number of frames) | No M<sup>+</sup><br>ions | K<sup>+</sup> | Na<sup>+</sup> | NH<sub>4</sub><sup>+</sup> |
| --- | --- | --- | --- | --- |
| No M<sup>+</sup> ions<br>(267) | N/A | 10<sup>-16</sup> | 10<sup>-15</sup> | 10<sup>-17</sup> |
| K<sup>+</sup><br>(198) | 10<sup>-16</sup> | N/A | 0.94* | 0.72* |
| Na<sup>+</sup><br>(192) | 10<sup>-15</sup> | 0.94* | N/A | 0.68* |
| NH<sub>4</sub><sup>+</sup><br>(190) | 10<sup>-17</sup> | 0.72* | 0.68* | N/A |

The null hypothesis was that the Pβ-O-P $\gamma$ angles in the $\alpha\beta\gamma$ -coordinated Mg-ATP complexes with different M<sup>+</sup> ions added result from normal distributions with equal mean values.
\* The null hypothesis was NOT rejected, no significant difference between samples

**Table S12.**
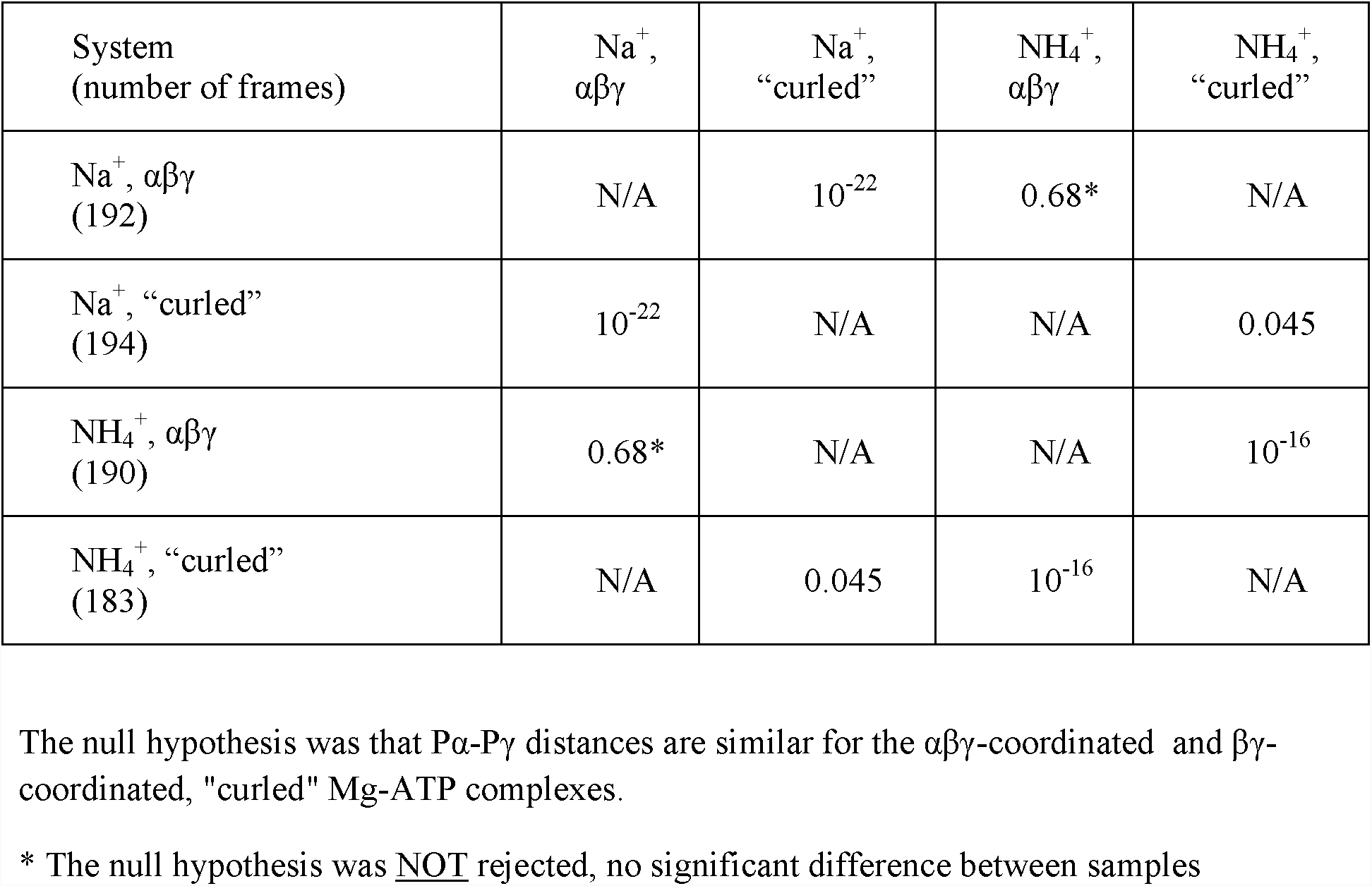
Comparison of the Pα-Pγ distance measurements for the αβγ-coordinated and “curled” βγ-coordinated Mg-ATP complexes

